# DNA-methylation markers associated with lung function at birth and childhood reveal early life programming of inflammatory pathways

**DOI:** 10.1101/2025.05.12.653131

**Authors:** Priyadarshini Kachroo, Katherine H. Shutta, Enrico Maiorino, Matthew Moll, Julian Hecker, Vincent Carey, Michael J. McGeachie, Augusto A. Litonjua, Juan C. Celedón, National Heart, Lung, and Blood Institute Trans-Omics for Precision Medicine (TOPMed) Consortium, Scott T. Weiss, Dawn L. DeMeo

## Abstract

**Rationale:** Lung function deficits may be caused by early life epigenetic programming. Early childhood studies are necessary to understand life-course trends in lung diseases.

**Objectives:** We aimed to examine whether DNA-methylation at birth and childhood is associated with lung function growth.

**Methods:** We measured DNA-methylation in leukocytes from participants in two childhood asthma cohorts (CAMP [n=703, mean-age 12.9 years] and GACRS [n=788, mean-age 9.3 years]) and cord blood from participants in the VDAART study (n=572) to identify CpGs and pathways associated with lung function.

**Results:** We identified 1,049 consistent differentially methylated CpGs (608 relatively hypermethylated) across all three studies (FDR-P<0.05). Relatively hypomethylated CpGs were enriched for gluconeogenesis, cell adhesion and VEGF signaling. Relatively hypermethylated CpGs were enriched for Hippo, B-cell and growth hormone receptor signaling. Functional enrichment suggested potential regulatory roles for active enhancers and histone modifications. Additionally, enrichment in PI3K/AKT and Notch pathways in males and enrichment in hormonal pathways in females was identified. Gaussian graphical models identified sex-differential DNA-methylation nodes and hub scores at birth and childhood. Integrating with previously identified polygenic risk scores for asthma and drug-target enrichment identified seven robust genes including *MPO*, *CHCHD3, CACNA1S, PI4KA, EP400, CREBBP* and *KCNA10* with known associations as biomarkers for asthma severity and drug targets for airway inflammation.

**Conclusions:** Epigenetic variability from birth through puberty provides mechanistic insights into fetal programming of developmental and immune pathways associated with lung function. These early life observations reveal potential targets for mitigating risk for lung function decline and asthma progression in later life.

**Key messages:**

- We identified consistent DNA methylation signatures between birth and childhood in critical metabolic, lung development and immune pathways that were associated with lung function and may be influenced by sex and genetics.
- Our integrative findings provide a deeper understanding for accelerated lung function decline across the life-course and could pave the way for translational interventions for lung diseases based on epigenetic plasticity.

## INTRODUCTION

Lung function (LF)(1) impairment often has origins in early life(2) and is influenced by a combination of genetics, prenatal or postnatal exposures, and environmental factors. Life-long consequences may include an early onset of chronic lung diseases including asthma and Chronic Obstructive Pulmonary Disease (COPD)(3) which share clinical features but also demonstrate substantial heterogeneity(4) and impart a major global health burden(5–9).

Asthma progression is associated with reduced lung growth and an accelerated rate of decline in FEV_1_ and FEV_1_/FVC(10). Children with reduced baseline FEV_1_ may be at risk of exacerbations, fixed airflow obstruction and early COPD(11–13). While several genetic variants may influence an individual’s asthma risk(14), LF genetic risk loci account for a fraction of the overall heritability with modest effects(12, 15, 16). Epigenetic marks including DNA methylation (DNA-m) are influenced by genetics and environment(17) and play an important role in lung development(18).

Prior studies have robustly associated DNA-m(19) with LF(20–24) but large-scale studies are needed to comprehensively explore their relationship from early life. A large-scale meta-analysis by Lee *et al*. expanded our knowledge on ancestry-specific epigenetic associations with LF, but their data mostly included older adults and sex differences were not considered (25). DNA-m levels are particularly impacted by sex during adolescence, and this may further play a role in sex-specific risks of respiratory conditions across the life course(26–28). Previously, DNA-m studies of pre-adolescent participants identified CpGs associated with sex-specific LF trajectories (age 10, 18 and 26 years)(27). Using the same cohorts, sex-specific DNA-m and gene expression patterns were identified at birth that correlated with LF in adolescence (29). Growing evidence suggests that investigating CpGs associated with high-risk LF trajectories (27, 30) may have the potential to identify specific inflammatory markers in relation to distinct asthma heterogeneity and inform preventative interventions for early-onset COPD.

This study examined epigenome-wide associations with multiple LF outcomes in three cohorts: the Genetic Epidemiology of Asthma in Costa Rica Study (GACRS), the Childhood Asthma Management Program (CAMP) and the Vitamin D Antenatal Asthma Reduction Trial (VDAART). LF trajectories associated with COPD were additionally evaluated in CAMP and asthma outcomes at age 6 years were evaluated in VDAART; sex-stratified associations were also assessed. We integrated polygenic risk scores for asthma to capture robust DNA-m signatures independent of genetic risk for asthma. Finally, we used several integrative epigenomic tools to ascertain potential biomarkers adding functional relevance for respiratory disease biology.

## METHODS

### Overall Study Design

The overall goal of this study was to identify early life predictors of LF in childhood asthma using genome-wide DNA-m from diverse cohorts (**Figure 1**). We utilized two childhood asthma studies for discovery (CAMP, n=703) and replication (GACRS, n=788) to identify CpGs associated with LF. To test an early origins hypothesis, we further investigated associations in the umbilical cord blood DNA-methylome in association with LF at ages 5 and 6 years in VDAART (n=572). We then meta-analyzed our findings to allow a more comprehensive assessment of the DNA-m associations followed by several downstream integrative analyses (**Figure 1)**.

**Figure 1.**
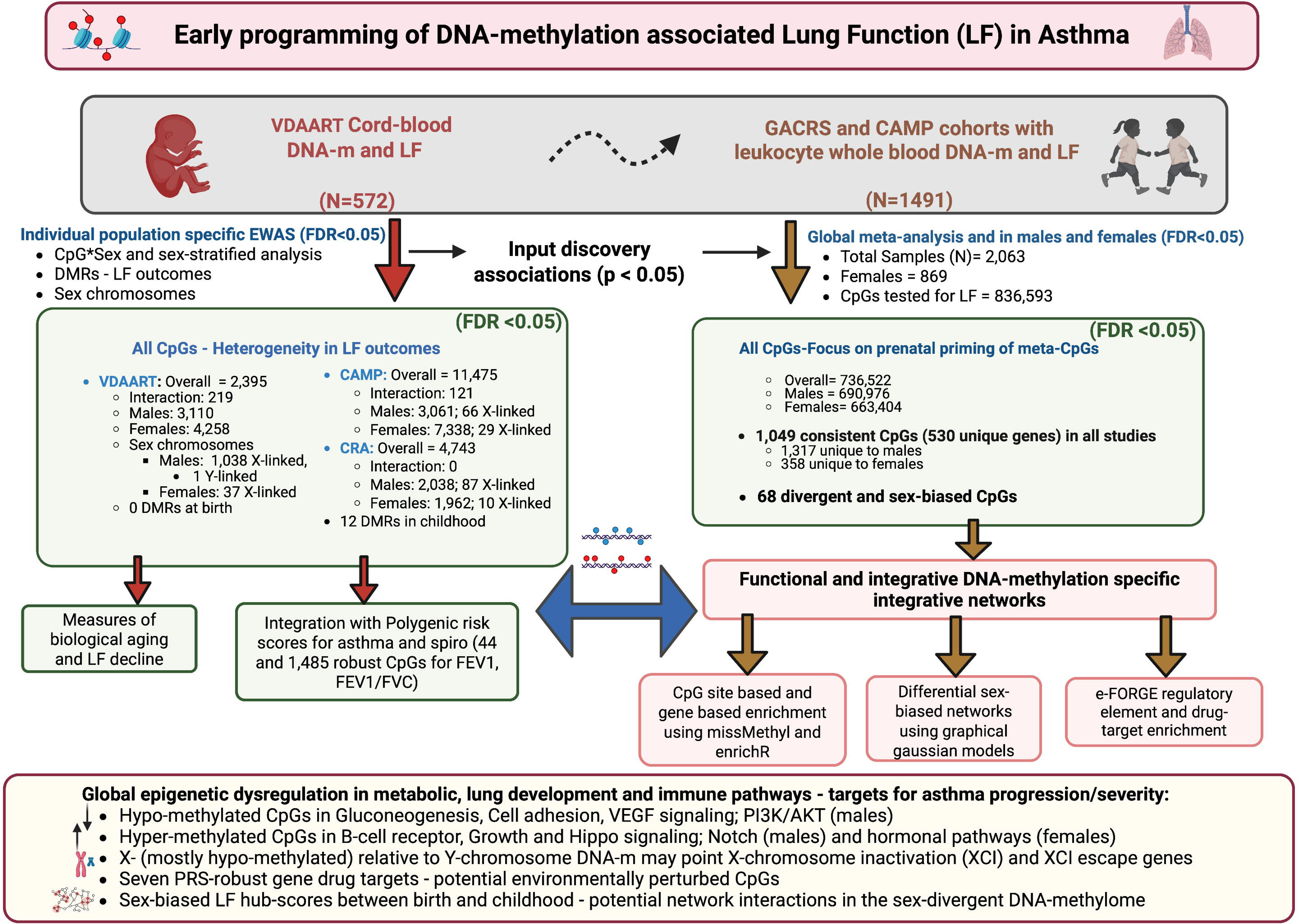
Overview of the conceptual framework and study design for the EWAS associations with LF and asthma outcomes in three different studies – CAMP, GACRS and VDAART. Created in BioRender. *Kachroo, P*. (2025) https://BioRender.com/502otie

In addition to commonly analyzed LF phenotypes such as Forced Expiratory Volume in 1 second, (FEV_1_,), Forced Vital Capacity (FVC) and FEV_1_/FVC, we evaluated the ratio of forced expiratory flow in the mid-portion of vital capacity divided by FVC (FEF_25-75_/FVC), as a surrogate measure of airway size relative to lung size, an understudied marker of dysanapsis previously associated with LF decline(31, 32). Longitudinal LF trajectories from CAMP(12, 13) were previously defined: Normal Growth (NG), deviations from the canonical NG pattern that can appear as either reduced growth (RG), early decline (ED) or a combination of RG and ED (any RG/any ED). Further, The Global Initiative for Obstructive Lung Disease (GOLD) stages for COPD were used for CAMP subjects who met the criteria for LF impairment at their last spirometry visit (aged 23 to 30 years)(13).’

Written informed consent and assent were obtained from both the parents and the participating children. Details on study populations and spirometry assessments are provided in the online supplement.

### Statistical analyses

#### Association of DNA-methylation with asthma and LF phenotypes

Data preprocessing and quality control methods are detailed in the Online Supplement. Mainly, to analyze DNA-m (hg19 reference genome), we used logit transformed β-values (M-values approximated by log2(β /(1-β)) (31). A multivariable robust regression model implemented in the robustbase R package(32) was used to analyze DNA-m M-values as predictor for LF/asthma outcomes adjusting for known covariates (see online supplement). DMRCate(33) was used to identify differentially methylated regions (DMRs).

To enrich insights into developmental origins, we further performed a meta-analysis using individual study-specific and male- and female-associated differentially-methylated positions (discovery DMPs, P-value<0.05) using inverse variance-weighted fixed-effects models implemented in the METAL software(34). The DMPs with a consistent direction of effect in all three or in at least two of the three studies (FDR<0.05) were retained for downstream analyses.

#### Functional downstream and integrative epigenomics

To assess whether the significant LF-associated DMPs were attenuated by genetic risk, we performed adjustment for polygenic risk scores for asthma (PRS_asthma_)(35) and a composite spirometry-based COPD PRS (FEV_1_, FEV_1_/FVC; PRS_spiro(36)_), both which were developed in external training cohorts(37, 38) and calculated in CAMP. We also tested models including the main effects and interaction terms for PRS X CpG. CpG site-based gene ontology and pathway enrichment analysis using KEGG were performed using missMethyl(39). The terms with ≥2 genes at a FDR-P<0.05(40) were regarded as significantly enriched. To get the most functionally relevant network representation, an integrative gene-based enrichment and visualization was performed for the meta-analyzed and male- or female-specific DMPs against five gene-set libraries (Gene Ontology, KEGG, GWAS Catalog, DisGeNET) using the knowledge-graph database and web-server enrichr-KG (https://maayanlab.cloud/enrichr-kg)(41). Further, for the LF-associated CpG DMPs that exist in both males and females but in opposite direction of effect, Gaussian graphical models (GGMs) were constructed to identify differential CpG nodes and hub scores specific to sex-stratified male-female network modules (details in Online Supplement) using the CRAN package huge(42).

Using the experimentally-derived Functional element Overlap analysis of ReGions from EWAS (eFORGE)(43, 44) integrative epigenomics tool, we further explored whether our meta-analyzed LF–associated DMPs were enriched in regulatory elements from the Roadmap Epigenomics Mapping Consortium across more than 20 cell/tissue types. We also applied the drug perturbation gene set enrichment analysis (dpGSEA)(45) (https://github.com/sxf296/drug_targeting) to the CpG-mapped genes and the directionality of effect of their DMPs and identified phenotypically relevant approved or experimental drug targets of clinical relevance derived using the Broad Institute’s Connectivity Map (CMAP) and the Library of Integrated Network-based Cellular Signatures (LINCS) framework. The top 50 drug-targeted genes with the cell-line information were evaluated. The gene-drug targets that showed a significant enrichment as well as target compatibility score at both 90% and 95% confidence interval from both databases CMAP and LINCS(46), were retained.

Biological age may reflect distinct developmental perturbations at birth and during childhood(47, 48). Therefore, we also applied generalized linear models to examine the association between the LF outcomes and measures of epigenetic age acceleration using the R/Bioconductor package methylclock(49). Elastic Net (EN) clock(50) trained on childhood data was applied for CAMP and GACRS and the EPIC clock(51) trained for gestational age predictions was applied for VDAART.

## RESULTS

The basic characteristics of each study population have been provided (**Table 1** and detailed in **Suppl. Tables E1-3**. Phenotypic data were available for 703 CAMP participants, 788 GACRS participants and 352 VDAART participants.

**Table 1.**
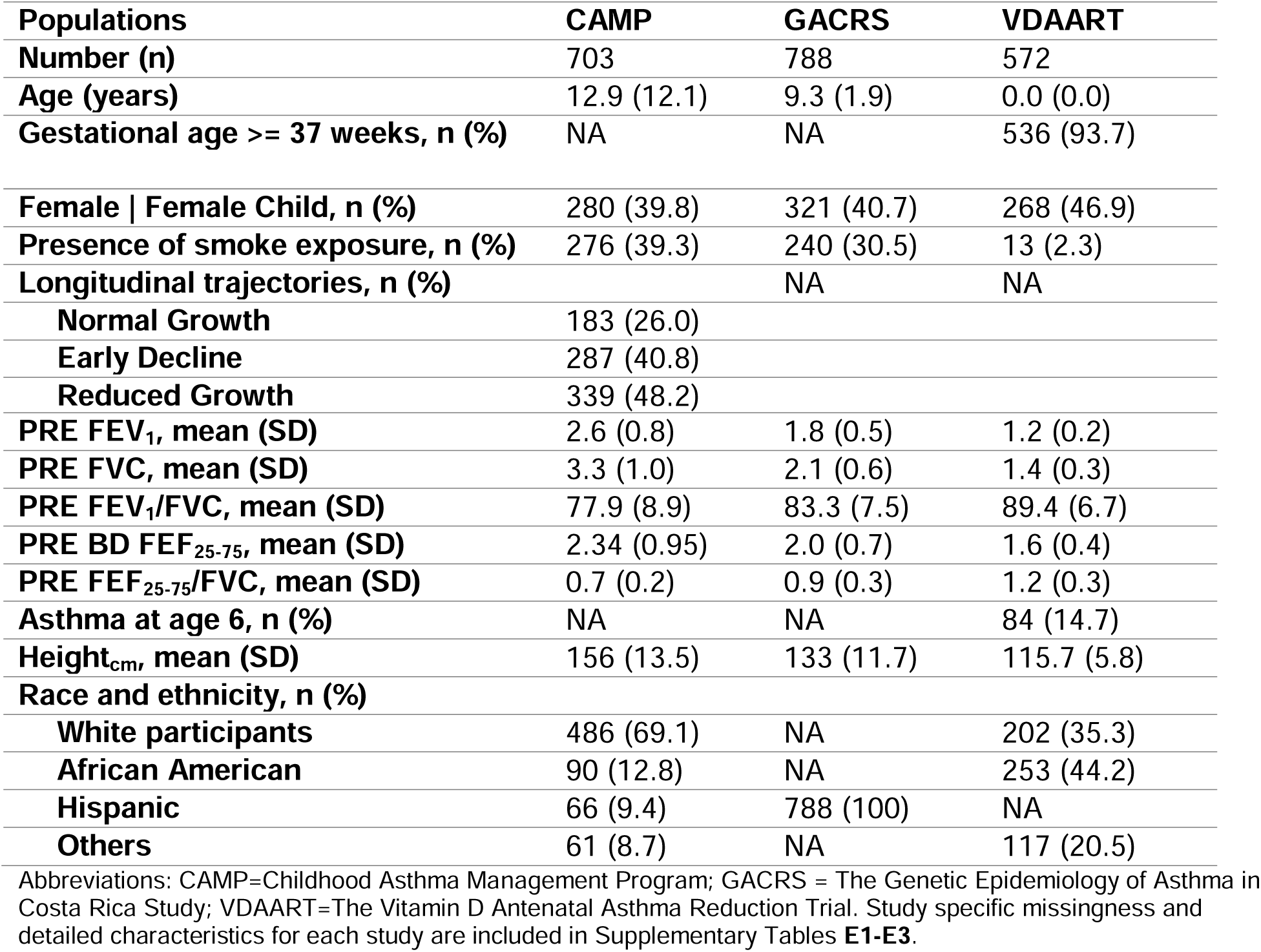
Characteristics of the study samples and participants in all three cohorts with data on DNA-methylation and lung function outcomes (N=2,063)

### Associations between DNA-methylation at birth/childhood and LF

We identified several LF-associated DMPs in the VDAART birth cohort (cord-blood) (**Table E4, Figure E1**). Between the two childhood cohorts GECRA and CAMP, 803 overlapping CpG DMPs were associated with either FEV_1_/FVC, FEF_25-75_ and FEF_25-75_/FVC, with 99.8% in consistent direction of effect (**Tables E5-E7**, detailed in Online Supplement). Fewer associations replicated between cord-blood and childhood (**Figure E2, Table E8, see online supplement**). Nominally significant associations (P<0.05, **Table E4**, **Figure E2-E3**) were identified for the RG and ED trajectories in CAMP and for asthma development in VDAART (see Online Supplement).

Based on the exact CpG coordinates (chromosome, start and end) and a stringent threshold of ≥4 CpGs within the associated region, we identified 11 DMRs for FEV_1_/FVC (mapping to 11 genes [*PCYT1A, IL4, EPX, EVL, RASSF2, VTI1A, TLDC2, FBXO7, IL5RA, IGF1R, HS2ST1*]) and one significant DMR for FEF_25-75_/FVC (mapping to *URI1*) that were replicated between GACRS and CAMP (**Table E9**). We did not identify any LF-associated DMRs in VDAART (FDR<0.05).

Analyzing LF-associated DNA-m in the sex chromosomes identified mostly X-chromosome associated differential methylation between males and females. Relatively fewer LF-associated X-chromosome DMPs (n=37) were identified in females, that were globally hypo-methylated in gene body and promoter-associated regions compared to male X-chromosome across all studies; 16 DMPs had increased DNA-methylation levels. In males, 652 X-chromosome DMPs were hyper-methylated and 172 DMPs were hypo-methylated (**Table E10**).

Epigenetic age acceleration was significantly associated with reduced LF in CAMP; cord blood age acceleration was associated with an increased LF in VDAART by age 5-6 (**Table E11,** see Online Supplement).

### Meta-analysis – Recapitulation of LF-associated DMPs with adult COPD

A total of 9,851 meta-DMPs were shared between all LF outcomes (**Figure 2**). We identified 1,049 (812 unique) meta-LF DMPs (**Table 2**, FDR<0.05) having consistent direction of effect across all studies, with 338 hypo-(209 genes) and 474 (326 genes) hyper-methylated (**Table E12)**; 47 DMPs were associated with at least three LF traits (**Figure 3A, Table E12**); the top five consistent DMPs are highlighted in **Table 3**. Three of the 1,049 meta-DMPs were also associated with the reduced lung growth-trajectory DMPs (cg20981347, cg26657392: *MFSD12* and cg17950165: *LINC01182*). Nine meta-DMPs (flagged in **Table E12**) were shared across seven blood EWAS studies in adults(52) in association with COPD (cg04637264:*KCNIP2*, cg09598552:*CDH23, cg09646173:PDE6A*), FEV_1_ (cg09646173:*PDE6A*, cg12077460:*MFHAS1*) and FEV_1_/FVC (cg12147622, cg00762550:*LIG3*, cg01878963:*RAP1A*, cg16518176:*PES1P1*) DMPs, while cg00278366:*RAD9B* was common to all.

**Figure 2.**
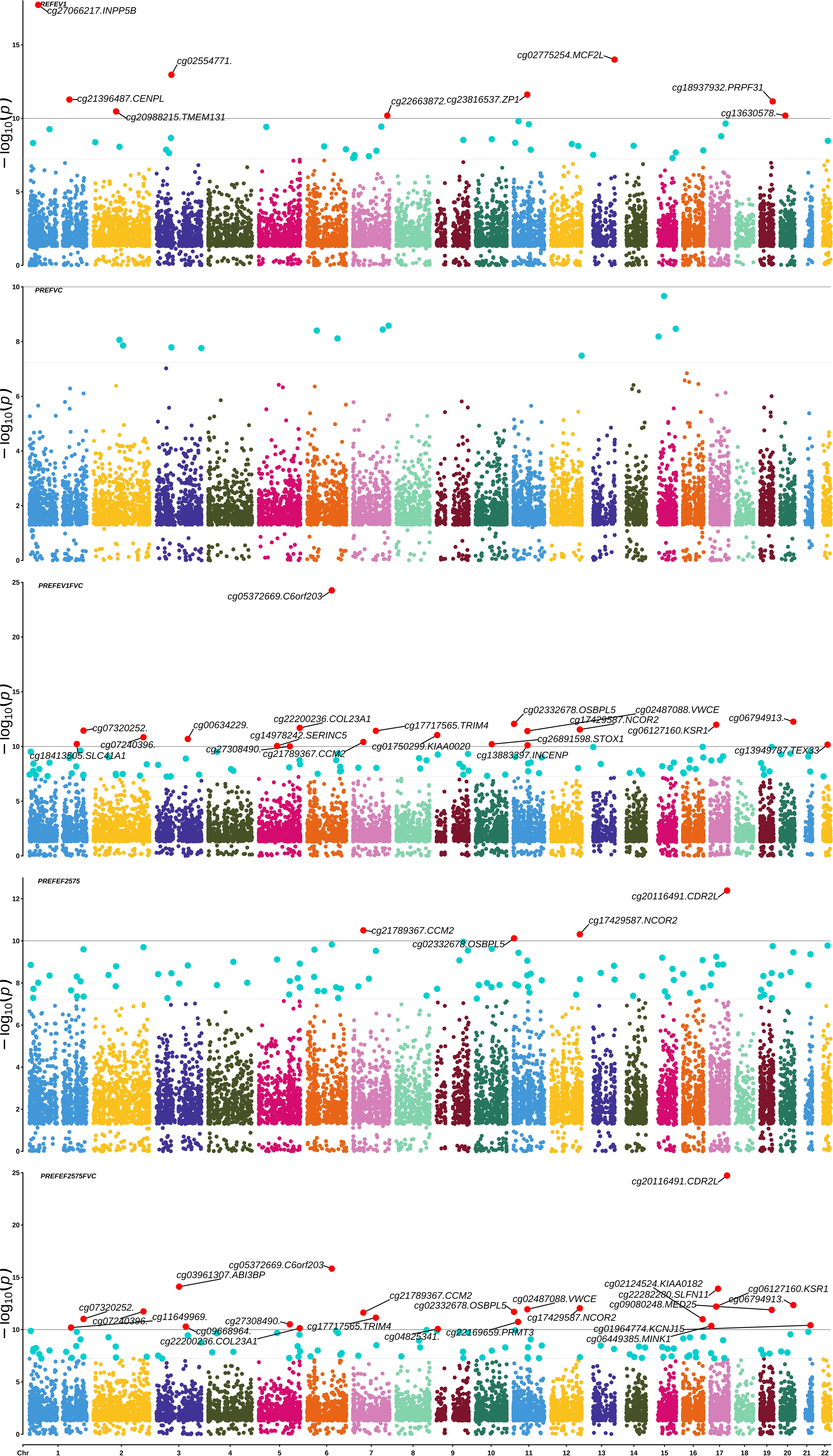
Multi-trait PheWAS graphical representation of the manhattan plot for the meta-analyzed EWAS associations. Only the overlapping DMPs across all tested LF phenotypes were used as input for this plot. Highlighted associations in blue are at a genome-wide threshold (p-value threshold=5.8x10^-8^). Highlighted associations in red are at a p-value threshold of 1x10^-10^ for better visualization of the top hits overlapping across multiple phenotypes. The density of CpGs on each chromosome are shown by the legend key with lowest to highest density ranging from green to yellow to red color.

**Figure 3.**
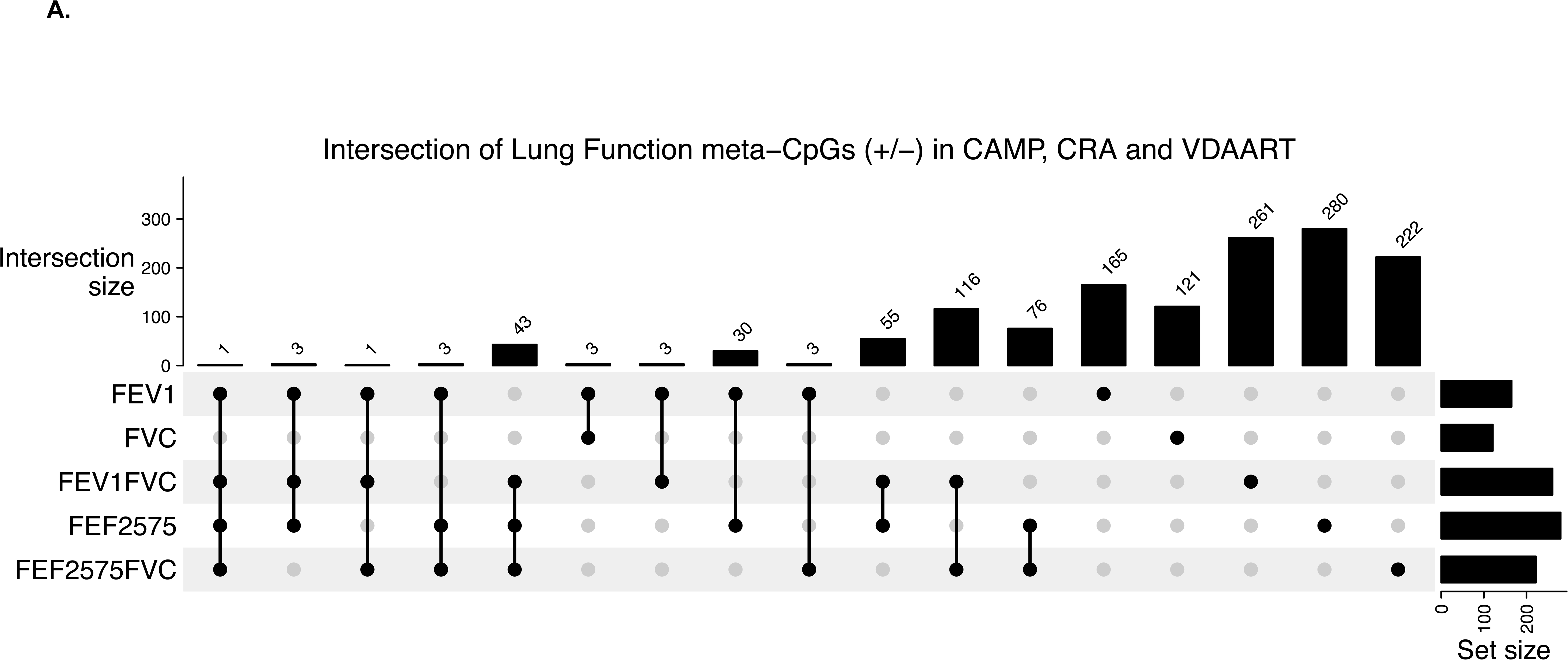
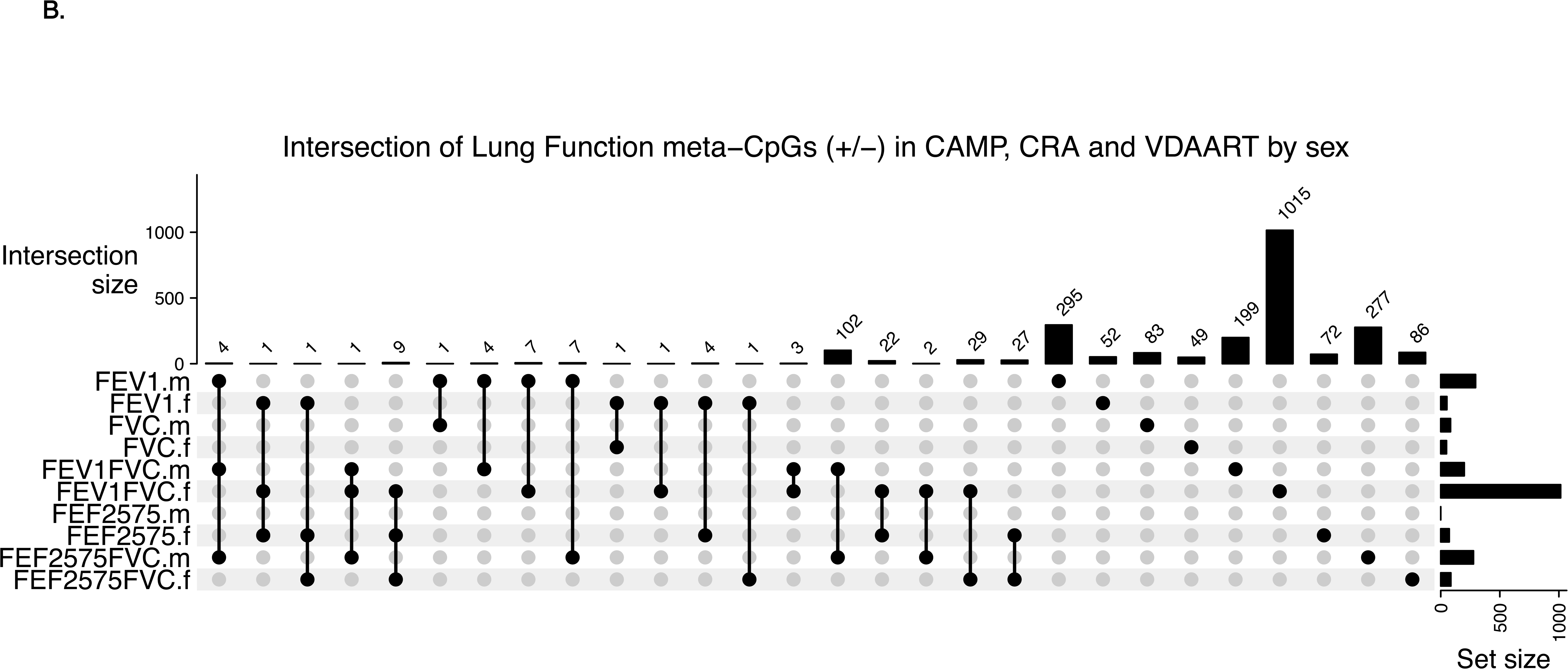
Upset plot showing the Intersection of differentially methylated CpGs from the meta-analysis in consistent direction of DNA-methylation effect across all three cohorts at FDR < 0.05 **A.** across LF phenotypes **B.** across LF phenotypes in males and females.

**Table 2.** Statistics and Number of epigenome-wide differentially methylated associations from the meta-analyses across three independent study populations: GACRS, CAMP and VDAART.

| **Meta-analyzed phenotype across all three studies** | **FEV<sub>1</sub>** | **FVC** | **FEV<sub>1</sub>/FVC** | **FEF<sub>25-75</sub>** | **FEF<sub>25-75</sub>/FVC** |
| --- | --- | --- | --- | --- | --- |
| **Associations tested** |  |  |  |  |  |
| Overall | 132,109 | 194,765 | 183,255 | 152,355 | 174,109 |
| Males | 157,142 | 157,759 | 160,359 | 159,318 | 159,659 |
| Females | 139,063 | 147,010 | 166,568 | 153,720 | 166,407 |
| **Associations at FDR<0.05** |  |  |  |  |  |
| Overall | 110,919 | 170,309 | 165,236 | 134,015 | 156,043 |
| Males | 136,910 | 132,933 | 141,759 | 138,233 | 141,141 |
| Females | 116,869 | 120,345 | 146,116 | 133,460 | 146,614 |
| **Consistent at FDR<0.05** |  |  |  |  |  |
| Overall | 165 | 121 | 261 | 280 | 222 |
| Males | 295 | 83 | 199 | 463 | 277 |
| Females | 52 | 49 | 99 | 72 | 86 |

**Table 3.** Top five CpGs with available gene annotations and showing consistent DNA-methylation associations with LF outcomes meta-analyzed between GACRS, CAMP and VDAART cohorts represented by the ‘Direction’ column respectively (FDR < 0.05). The list is sorted by FDR.

| Phenotype | CpG probe | chr | pos | Gene | Effect | Direction | P.value | FDR | Relation to Island | Gene Context |
| --- | --- | --- | --- | --- | --- | --- | --- | --- | --- | --- |
| FEV<sub>1</sub> | cg12684668 | chr5 | 150,403,466 | *GPX3* | -<br>0.0903 | --- | 7.56E-08 | 8.76E-05 | S_Shelf | Body |
|  | cg12233571 | chr5 | 36,613,509 | *SLC1A3* | 0.0732 | +++ | 1.77E-07 | 0.00016674 | OpenSea | Body |
|  | cg01978458 | chr7 | 2,683,257 | *TTYH3* | 0.0894 | +++ | 3.11E-07 | 0.00024335 | N_Shelf | Body |
|  | cg07447194 | chr3 | 148,711,966 | *GYG1* | 0.0596 | +++ | 3.66E-07 | 0.0002705 | S_Shore | Body |
|  | cg12120947 | chr6 | 32,166,652 | *NOTCH4* | 0.0594 | +++ | 3.71E-07 | 0.0002705 | S_Shelf | Body |
| FVC | cg01900413 | chr11 | 128,419,356 | *ETS1* | 0.1068 | +++ | 1.44E-06 | 0.00096312 | Island | Body |
|  | cg13618478 | chr19 | 56,549,544 | *NLRP5* | -<br>0.0751 | --- | 1.54E-06 | 0.00100793 | OpenSea | ExonBnd |
|  | cg21817833 | chr14 | 52,118,115 | *FRMD6* | 0.1585 | +++ | 1.62E-06 | 0.00103473 | Island | TSS1500 |
|  | cg09405702 | chr8 | 12,588,030 | *LONRF1* | 0.049 | +++ | 1.79E-06 | 0.00109532 | OpenSea | Body |
|  | cg07167860 | chr13 | 93,136,776 | *GPC5* | 0.0411 | +++ | 2.58E-06 | 0.00130417 | OpenSea | Body |
| FEV<sub>1</sub>/FVC | cg25627789 | chr7 | 132,694,521 | *CHCHD3* | 3.5519 | +++ | 1.16E-12 | 3.86E-09 | OpenSea | Body |
|  | cg14978242 | chr5 | 79,501,131 | *SERINC5* | 2.2488 | +++ | 9.02E-11 | 7.21E-08 | OpenSea | Body |
|  | cg04141008 | chr16 | 3,843,232 | *CREBBP* | 3.7667 | +++ | 2.95E-10 | 1.73E-07 | OpenSea | Body |
|  | cg05380077 | chr11 | 63,272,225 | *LGALS12* | 3.6992 | +++ | 5.91E-10 | 2.82E-07 | OpenSea | TSS1500 |
|  | cg18460265 | chr1 | 111,062,669 | *KCNA10* | 2.7615 | +++ | 1.09E-09 | 4.39E-07 | OpenSea | TSS1500 |
| FEF<sub>25-75</sub> | cg16107105 | chr7 | 150,646,704 | *KCNH2* | 0.1672 | +++ | 1.32E-10 | 2.89E-07 | N_Shore | Body |
|  | cg09845476 | chr9 | 127,113,228 | *NEK6* | 0.2199 | +++ | 1.63E-10 | 3.25E-07 | OpenSea | 3'UTR |
|  | cg26054828 | chr10 | 75,619,500 | *CAMK2G* | 0.37 | +++ | 1.82E-10 | 3.33E-07 | OpenSea | Body |
|  | cg12120947 | chr6 | 32,166,652 | *NOTCH4* | 0.163 | +++ | 2.60E-10 | 3.94E-07 | S_Shelf | Body |
|  | cg18337287 | chr19 | 930,871 | *ARID3A* | 0.1486 | +++ | 3.72E-10 | 4.68E-07 | N_Shore | Body |
| FEF<sub>25-75</sub>/FVC | cg13774539 | chr2 | 74,612,706 | *LOC100189589* | 0.1213 | +++ | 1.02E-10 | 5.82E-08 | OpenSea | TSS200 |
|  | cg14978242 | chr5 | 79,501,131 | *SERINC5* | 0.074 | +++ | 2.11E-10 | 9.96E-08 | OpenSea | Body |
|  | cg18181035 | chr3 | 128,214,460 | *GATA2-AS1* | 0.1578 | +++ | 8.46E-10 | 2.87E-07 | N_Shore | Body |
|  | cg21271570 | chr13 | 44,961,267 | *SERP2* | 0.0897 | +++ | 3.24E-09 | 8.53E-07 | OpenSea | Body |
|  | cg12242115 | chr12 | 122,251,166 | *SETD1B* | -<br>0.1273 | --- | 3.78E-09 | 9.66E-07 | S_Shore | Body |

**Table 4.** Gene-drug targets with significant enrichment and target compatibility score for the meta-analyzed CpG associations specific to each LF outcome.

| PHENOTYPE | CMAP database | LINCS database | OVERLAP from both databases | Number of Drug Targets in CMAP* | Number of Drug Targets in LINCS* |
| --- | --- | --- | --- | --- | --- |
| FEV<sub>1</sub> | 2 (*GPX3*, *SLC1A3*) | 6 (*GYG1*, *NOTCH4*, *MPO*, *ABAT*, *GPX3*, *SLC1A3*) | *GPX3*<br>*SLC1A3* | 2<br>6 | 12<br>35 |
| FVC | 2 (*ETS1*, *LONRF1*) | 1 (*ETS1*) | *ETS1* | 4 | 56 |
| FEV<sub>1</sub>/FVC | 8 (*CACNA1S*, *SLC38A10*, *PPM1H*, *TINAGL1*, *CHCHD3*, *SERINC5*, *KCNA10*, *EP400*) | 7 (*CREBBP*, *PI4KA*, *RPS6KC1*, *CHCHD3*, *SERINC5*, *KCNA10*, *EP400*) | *CHCHD3*<br>*SERINC5*<br>*KCNA10*<br>*EP400* | 2<br>1<br>2<br>1 | 15<br>74<br>2<br>22 |
| FEF<sub>25-75</sub> | 5 (*CAMK2G*, *NOTCH4*, *ARID3A*, *SERINC5*, *SDC3*) | 11 (*KCNH2*, *ARID3A*, *LTBP1*, *RAB20*, *SIGLEC8*, *ITSN1*, *CAMK2G*, *NOTCH4*, *ARID3A*, *SERINC5*, *SDC3*) | *CAMK2G*<br>*NOTCH4*<br>*ARID3A*<br>*SERINC5*<br>*SDC3* | 1<br>4<br>2<br>2<br>6 | 18<br>9<br>11<br>73<br>23 |
| FEF<sub>25-75</sub>/FVC | 6 (*SERINC5*, *SETD1B*, *RAB20*, *KCNA10*, *EP400*, *CACNA1S*) | 8 (*KSR1*, *SGMS1*, *CANX*, *SERINC5*, *SETD1B*, *RAB20*, *KCNA10*, *EP400*) | *SERINC5*<br>*SETD1B*<br>*RAB20*<br>*KCNA10*<br>*EP400* | 2<br>1<br>1<br>2<br>1 | 90<br>14<br>27<br>2<br>21 |

\*See supplemental **Table E16** for drug targets

### Identification of PRS-robust CpGs after PRS adjustment

PRS integration reduced our sample size in CAMP by almost 38%, yet the regression coefficients remained unchanged for 44/387 DMPs that remained robustly associated with FEV_1_ (**Table E13, Figure E4A**). Notably, three of the 44 DMPs were associated in the meta-analysis for CpG-FEV_1_ associations: cg04266202 (*MPO)* and cg06070625 (*MITF*). The regression coefficients remained unchanged for 1,485/3,485 CpGs that remained robustly associated with FEV_1_/FVC (**Table E14, Figure E4B**). Notably, 10 were associated in the meta-analysis of CpG-FEV_1_/FVC associations (cg25627789:*CHCHD3*, cg12046819:*SGMS1*, cg00068153:*CACNA1S*, cg09662086:*PI4KA*, cg05486260:*FAM135B*, cg00764582:*EP400*, cg22330572:*AZIN1-AS1*, cg04141008:*CREBBP*, cg18460265: *KCNA10*). Similar trends were observed when evaluating LF-associated DMPs and PRS_spiro_ models with potential relevance for COPD risk (**Figure E4**).

### Integrative epigenomic analyses identified pathways with functional implications

Of the meta-analyzed LF-DMPs, hypo-methylated CpGs were enriched for growth-related and developmental pathways (**Table E15**) including Vascular Endothelial Growth Factor (VEGF) receptor signaling (*FGF9, PTK2, VAV2, VEGFC*), semaphorin-plexin signaling (*PLXND1, PLXNA4*), glucose metabolism (*PGM2, ADPGK, FBP2*) and negative Wnt signaling regulation (*SOX30, FGF9, LATS2, UBAC2, KREMEN1, NKD1*). Hyper-methylated CpGs were enriched for developmental and immune function related processes and pathways (**Table E15**) including positive regulation of B cell receptor/antigen receptor-mediated signaling (*PRKCH, SLC39A10, CD81, LGALS3*), insulin receptor signaling (*SORBS1, OSBPL8, PTPN1*), growth hormone receptor signaling (*PTPN1, STAT5A*), Wnt signaling (*CSNK1E, WNT3*), and the Hippo signaling pathway (*RASSF1, FRMD6, CSNK1E, RASSF4*). The gene-based enrichment identified similar and new integrative associations from KEGG, GWAS Catalog and DisGeNET (**Figure 4**).

**Figure 4.**
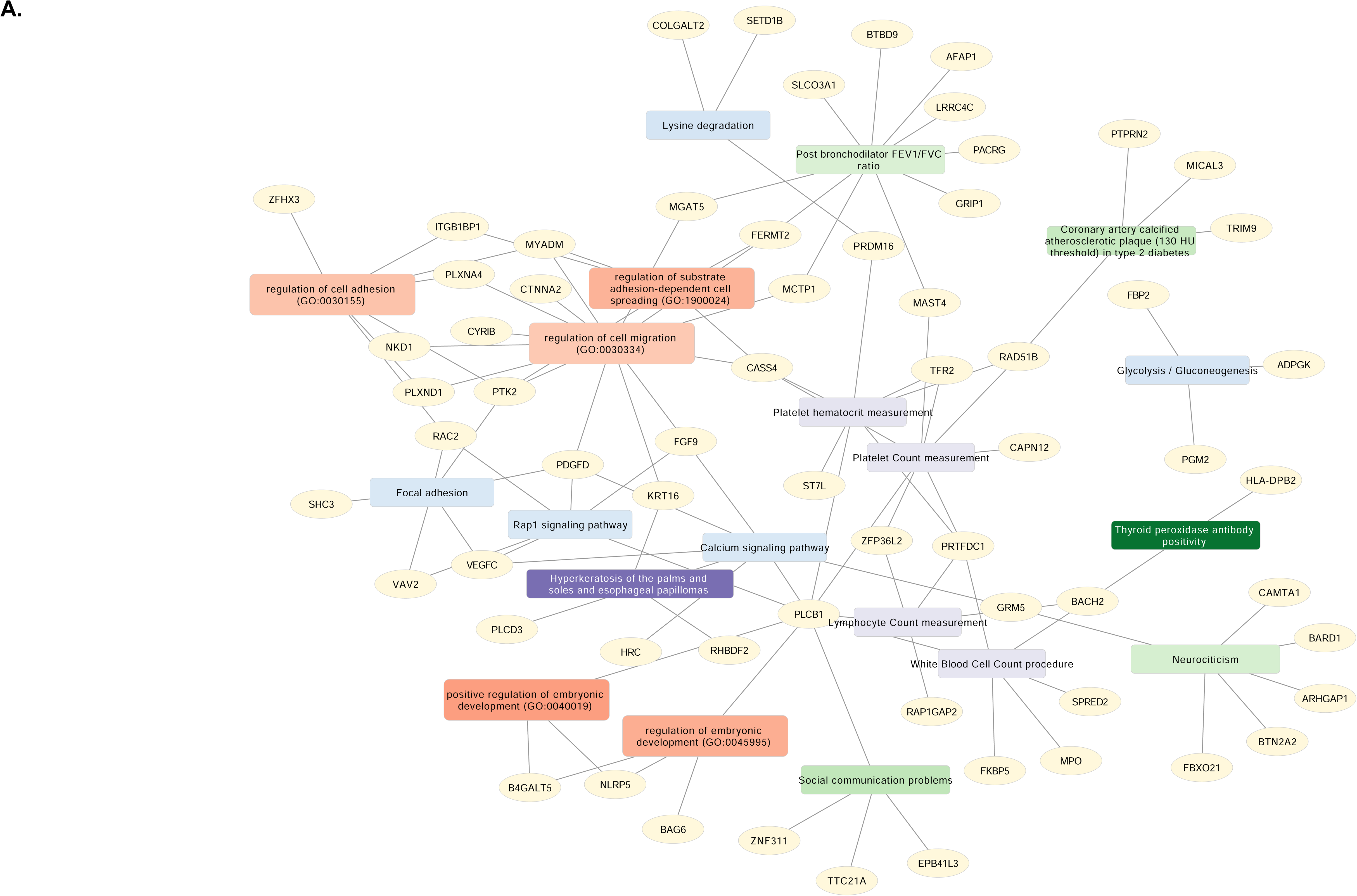
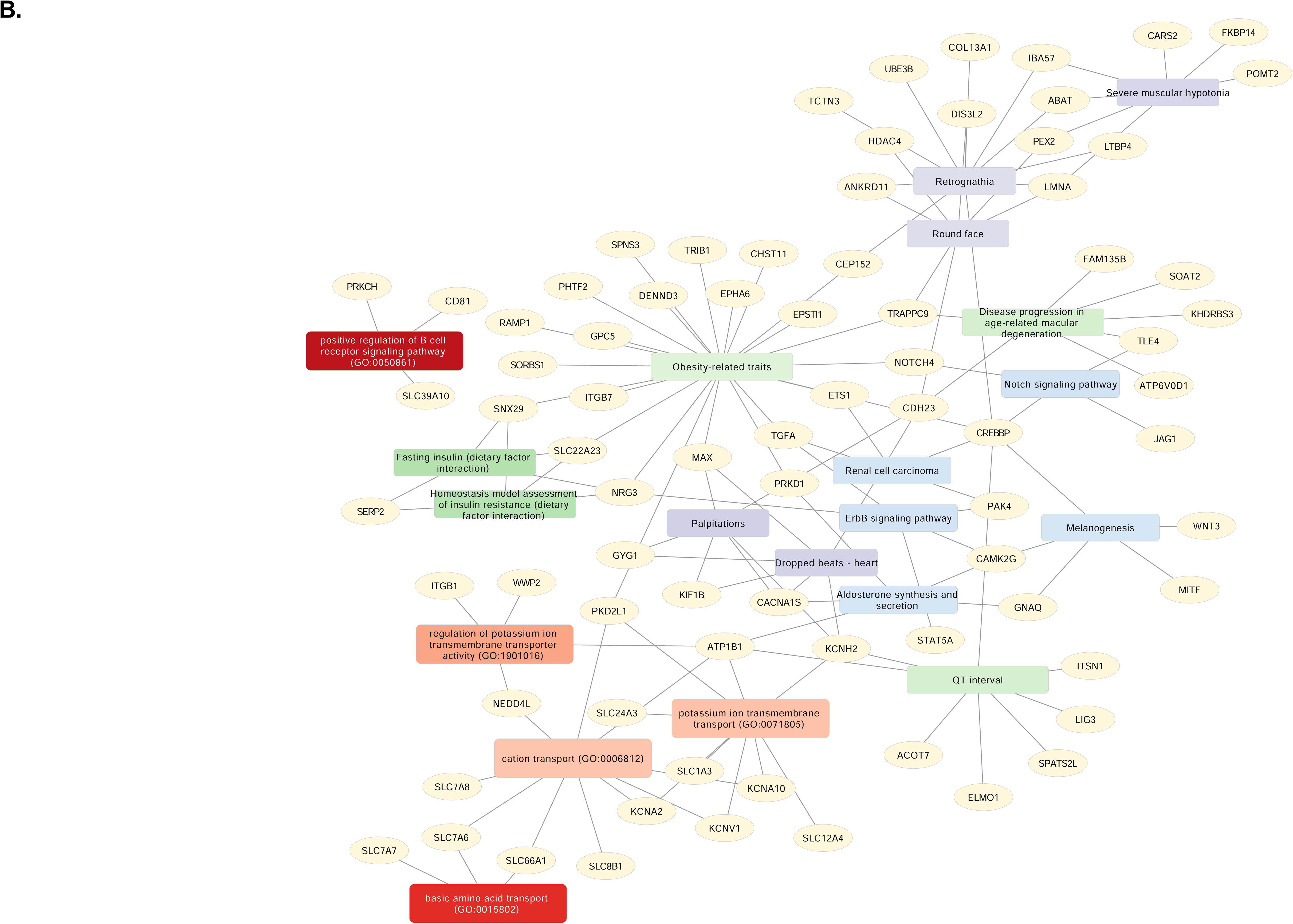
Functional integrative gene-based enrichment for genes annotated to the meta-analyzed hypo- and hyper-methylated CpGs consistent across all three studies and across all LF phenotypes: GACRS, CAMP and VDAART. Nodes are the genes; rectangles are the enriched terms. Each library source for the gene-based term enrichment is given a different color: Gene ontology (GO) Biological process (Red), KEGG (Blue), GWAS Catalog (Green) and DisGeNET (Purple), and the gradient is based on the z-score (the darker, the higher z-score). The hypo-methylated sub-network was enriched for biological processes: cell migration and adhesion, embryonic development; KEGG pathways: Calcium signaling, Glycolysis and Rap1 signaling; GWAS associations: post-bronchodilator FEV_1_/FVC ratio, type 2 diabetes, neurocognitive behaviors. The hyper-methylated sub-network was enriched for biological process: amino acid transport; KEGG pathways: B-cell receptor signaling, ErbB signaling, Aldosterone synthesis/secretion and Notch signaling pathway; GWAS associations: Obesity related traits, insulin resistance and age-related macular degeneration. **A.** Enrichment analysis for gene list based on hypo-methylated CpGs; **B.** Enrichment analysis for gene list based on hyper-methylated CpGs

Of all the identified phenotype-specific drug targets (**Table E16)**, 42 drug targets (19 unique genes) and 576 drug targets (27 unique genes) were identified using CMAP and LINCS respectively. Seven of those included the PRS-robust genes for FEV_1_ (*MPO*) and FEV_1_/FVC (*CHCHD3, CACNA1S, PI4KA, EP400, CREBBP, KCNA10*). Both databases identified 13 overlapping genes with several approved or known drug targets for respiratory and other health outcomes; *SERINC5, NOTCH4* and *EP400* had multi-phenotype associations (**Table 6**).

For further functional support of our findings, eFORGE integration for hypo-methylated CpGs showed enrichment of DNAase hotspots in the blood, lung and skin tissues, a strong enrichment of H3K4me1 in blood and transcriptionally active enhancers in several tissues including fetal lung tissue. The hyper-methylated CpGs showed enrichment of DNAase hotspots, a strong enrichment of H3K4me1 and H3K36me3 marks (**Figure E5**) and a weak transcription with enhancer activity across majority of the tissues including fetal lung.

### Sex stratified meta-analysis – Unique sex-specific and sex-divergent LF-DMPs

When stratified by sex, several consistent associations either for males (1,317 DMPs; 1,035 hyper-methylated) or for females (358 DMPs; 1 hyper-methylated) were identified between all studies (**Table 2, Table E17, Figure 3B**). Male-associated hypo-methylated DMPs were enriched for the PI3K-Akt signaling pathway and hyper-methylated DMPs were enriched for immune cell phenotypes, and Notch signaling. Female-associated hypo-methylated DMPs were enriched for GWAS genes for FEV_1_ and FEV_1_/FVC ratio, while the hyper-methylated DMPs were enriched for growth hormone, thyroid hormone, aldosterone and insulin signaling (**Table E18**).

Further, 76 (68 unique) DMPs exhibited opposite direction of effect between males and females (FDR < 0.05 in either males or females) and had consistent associations in at least two of the three studies for any LF outcome (**Table E17**). We constructed stratified GGMs on these DMPs and calculated hub scores of each node to assess how influential each DMP is within the female network and the male network (**Figure E6A**). This analysis yielded 11 DMPs (six with gene annotations) with more than 20% difference in hub scores between males and females from all three study populations (**Figure E6B**). One CpG annotated to *PLTP* gene (difference=0.22) was identified in GACRS, four CpGs annotated to *STX12* (difference=0.25), *LOC101928304* (difference=0.21), *KBTBD11* (difference=0.31) and *RCCD1* (difference=0.21) were identified in CAMP and 1 CpG annotated to *MAP3K7* (difference=0.36) was identified in VDAART. Additionally, eight DMPs were identified that were remarkably different between the birth cohort (VDAART) and either of the two childhood cohorts (GACRS, CAMP; **Figure E6C**).

## DISCUSSION

Epigenetic modifications capture pre- and post-natal environmental exposures, and these signatures may impact lung function and potentially lifelong susceptibility to chronic lung diseases like asthma and COPD(52). Prior large-scale EWAS studies(51) inadequately captured heterogeneity or were limited in scope, motivating us to comprehensively examine the relationship between DNA-m, lung function and asthma from birth to early adulthood. To our knowledge, this is the largest EWAS study of lung function in children at risk for and with asthma, revealing epigenetic dysregulation in key pathways including Hippo, Wnt and VEGF related signaling. *In silico* analyses have revealed gene-drug associations across three heterogeneous populations spanning birth and childhood and provide potential targets to explore further for primary prevention of obstructive lung disease.

Several population-specific LF-CpG associations replicated between GACRS and CAMP for FEV_1_/FVC and FEF_25-75_/FVC phenotypes, while many cord-blood associations were unique to VDAART, suggesting age-related heterogeneity in the epigenome. The LF-associated DMRs replicating between GACRS and CAMP included *IL4, EPX, IL5RA, IGF1R,* all previously associated with IgE-mediated respiratory diseases (53), allergic asthma(19) and reduced airway hyperresponsiveness in mice exposed to house-dust mite allergen(54). Such candidates could be of interest for epigenetic interventions focused on allergic asthma endotypes.

Our meta-analyzed LF-associated CpGs highlighted novel hypo- and hyper-methylated loci, pathway changes predicting LF decline, and metabolic pathway disruptions which may provide a clearer snapshot of the global DNA-m perturbations impacting asthma pathophysiology and COPD risk. VEGF signaling and glucose metabolism pathways suggest potential intervention targets. Particularly, upregulation of cell adhesion and VEGF signaling plays an important role in Th2 inflammation, regulating airway remodeling and hyper-responsiveness in both asthma and COPD(55). Previously, we identified enrichment of VEGFA-VEGFR2 signaling in fetal lung exposed to in-utero smoke(56) suggesting that some of these marks could be triggered by maternal exposures. Tissue-specific functional enrichment of H3K4me1 marks provided evidence of an additional regulatory role of active enhancers and histone modifications(57). Histone methylation disruption and increased VEGF driven by *IL13* has been linked to Th2 inflammation in asthma(58). Moreover, dysregulation in metabolic pathways may be observed during airway inflammation in respiratory diseases(59, 60). Experimental evidence further links VEGF and inhibition of glycolysis to hypoxia in pulmonary hypertension and endothelial dysfunction (61). Epigenetic perturbations in glucose metabolism suggests metabolic reprogramming in the genes mapping to this pathway(60), (62).

Hypermethylated CpGs associated with lung function were enriched for Hippo, B-cell and growth hormone receptor signaling pathways. Several prior studies have demonstrated that alterations in Wnt(63) and Hippo signaling(64) impact lung development and asthma progression(65). Previously, we have shown hyper-methylated CpGs in fetal lung exposed to in-utero smoke recapitulate in adult lung tissue and obstructive lung disease, with enrichment in Wnt and Hippo pathways(66). Balance between Wnt enhancers and inhibitors has demonstrated perturbations in airways from individuals with severe asthma(67). Hyper-methylation of LF-associated CpGs, such as *LATS2*, a regulator of Wnt-Hippo, can downregulate hippo signaling genes, which could be one potential mechanism to elevate Wnt signaling, disrupting the balance between Th2 and Th17 inflammatory responses in asthma. Positive regulation of B-cell receptor signaling was an enriched pathway for hyper-methylated CpGs, suggesting that B cell regulation could be an immune pathway dysregulated by epigenetic control in childhood and impact future risk for adult lung disease, given recent research supporting the role of the B cell in COPD pathogenesis(68). Genes mapping to the Insulin and growth hormone receptor signaling pathways could point us to the epigenetic regulation of metabolic syndrome affecting the uptake of glucose and lipids associated with asthma pathogenesis. In this pathway, the CpGs for *STAT5A* and *PTPN1* were hyper-methylated across the three studies and were around the transcription start site of the promoter region. Hyper-methylation of the *STAT5A* promoter has been associated with decreased expression in children with asthma(69). *PTPN1* promoter hyper-methylation was associated with type-2-diabetes(70). It is also noteworthy to find interactions between ErbB and aldosterone signaling in the hyper-methylated sub-network. Inhaled corticosteroids (ICS) are recommended to prevent asthma exacerbation in persistent asthma, and aberrant ErbB signaling mediates corticosteroid resistance driven by *IL13* in bronchial hyperresponsiveness(71). Further, ICS drugs mainly exert their effects on mineralocorticoids like aldosterone and glucocorticoid hormone receptors, therefore their prolonged exposure can cause defective DNA-binding or receptor mutations as seen in cortisol resistance(72); genes we have identified in the hypermethylated subnetwork may have pharmacoepigenetic relevance.

Existing studies show that LF-associated DNA-m in childhood and adolescence may vary by sex (73–75). We observed male enriched pathways included PI3K/AKT, linked to severe asthma outcomes, while female-specific enriched pathways involved thyroid, aldosterone, and insulin signaling. In contrast, sex-divergent methylation in specific genes may reveal insights into prenatal priming of age-related epigenetic perturbations associated with lung function and asthma in childhood. Exemplar genes with sex variable associations with lung function include *MAP3K7*, which is a known direct target for IL13 therapy in asthma(76, 77). Reduced *PLTP*, a phospholipid transport gene, which has been associated with neutrophil degranulation in COPD(78). *IL20RB*(*79*) and *TRH*(*80*) are critical lung maturation genes with strikingly different hub scores between birth and childhood; previously these genes have been linked with lung fibrosis development. Sex-specific DNA-m differences were also strikingly evident for genes on the X-chromosome, suggestive of potential X-chromosome mechanisms could impact lung function and potentially obstructive lung disease severity(81, 82). Our findings suggest that an in-depth sex-specific investigation should be the standard in large-scale studies. This would help characterize the role of epigenetic marks in sex-related features of lung function and prevalences of obstructive lung diseases across the life-course.

*In silico* approaches applied to our epigenetic associations identified key genes and potential clinical drug targets for further investigation and validation. *SERINC5* and *NOTCH4* were enriched for multiple LF phenotypes. Interestingly, *NOTCH4*, is regulated by Wnt-Hippo signaling(83), its inhibition can suppress Th17-mediated asthma hyper-responsiveness and vascular remodeling in COPD(84). For example, Maraviroc, the first approved CCR5 antagonist and Notch4 drug target, showed promise against severe respiratory conditions(85). *SERINC5* regulates viral infections through multiple immune pathways which could mitigate defective host defense mechanisms through targeting (86). Two of our drug targets also included PRS-robust genes *MPO*(*87*) and *CREBBP*(*88*), considered as biomarker for asthma severity and neutrophilic inflammation; their inhibition could attenuate oxidative damage.

Our study demonstrates that genetic signals may partially attenuate epigenetic signals. However, PRS adjustment did not fully attenuate CpGs associations suggesting these sites may be more susceptible to changes in DNA-m triggered by environmental exposures and potentially better targets for primary prevention through epigenetic plasticity. Only 20 (2.4%) of our 530 consistent meta-LF genes overlap with the lung function genetic associations identified by GWAS(37) suggesting low influence of genetic attenuation on these DNA-m signals. Likewise, Lee *et al*(25), reported limited overlap of their findings with genetic data. However, several of our identified meta LF-hits overlapped with the COPD and LF-associated CpGs across 17 previously published EWAS studies(52). Future studies should consider investigating causal effects on the epigenome and LF.

Our study has notable strengths in providing a broader understanding of birth and childhood DNA-m associations with lung function in asthma, using three multi-ethnic cohorts and various informatic methods to identify genes and pathways for future clinical study. Several limitations should also be acknowledged. Phenotypic variability, residual population substructure and small sample sizes in certain strata may account for lack of replication of some findings. Polygenic risk scores for asthma were only evaluated for CAMP, limiting replication of signals after adjustment for genetic risk. Further, we did not adjust for smoking-associated CpGs as a measure of second-hand smoke exposure, and this may reflect residual confounding in our findings.

In conclusion, we provide an integrative multidimensional framework demonstrating developmental plasticity and early-life programming of epigenetic mechanisms associated with lung function between birth and adolescence. Our findings inform the identification of genes and pathways for lung function in children with or at risk for asthma and highlight the epigenome as a next-generation target for therapeutic interventions for primary prevention of adult lung diseases and related inflammatory conditions.

## Acknowledgements

The authors thank all the participants, investigators and staff without whom this work could not have been accomplished. We gratefully acknowledge the individuals and studies that provided biological samples and data to TOPMed CAMP and GACRS and VDAART studies. We also appreciate the support of the NHLBI TOPMed initiative in facilitating the 850K EPIC array data generation using the Infinium® MethylationEPIC 850K BeadChip and contributing to overall research work. For a comprehensive list of TOPMed collaborators, refer to https://www.nhlbiwgs.org/topmed-banner-authorship

## Data sharing statement

All TOPMed data is person-sensitive, however it can be requested for access and can be made available through the TOPMed consortium after careful review and approval by the TOPMed Data Access Committee (https://topmed.nhlbi.nih.gov/). Participant consent and Data Use Limitations differs within and across TOPMed studies and should be requested individually. Additional documentation, such as of local IRB approval and/or letters of collaboration with the primary study PI(s) may be required.

The CAMP DNA methylation datasets analyzed in the current study are available at the database of Genotypes and Phenotypes (dbGaP) repository (phs001726. v2. p1) here: https://www.ncbi.nlm.nih.gov/projects/gap/cgi-bin/study.cgi?study_id=phs001726.v2.p1. The CRA DNA methylation datasets analyzed in the current study are available at the dbGaP repository (phs000988. v5. p1) here: https://www.ncbi.nlm.nih.gov/projects/gap/cgi-bin/study.cgi?study_id=phs000988.v5.p1.

All datasets used and/or analyzed during the current study could be requested from the corresponding author and contact study PIs and made available on reasonable request.

