## Supplementary material for "DNA-methylation markers associated with lung function at birth and childhood reveal early life programming of inflammatory pathways": Online Supplement

***Study populations:***

**CAMP and GACRS:**

The Childhood Asthma Management Program (CAMP) was a multi-center, double blind randomized clinical trial in 1041 children with mild-to-moderate asthma(1). The original 4.3-year trial randomized 1041 children 5 to 12 years of age from December 1993 to September 1995. The results of the clinical trial and follow-up phase have also been published(1-5). CAMP was approved by the Institutional Review Boards of Brigham and Women's Hospital and the other participating centers.

The Genetic Epidemiology of Asthma in Costa Rica Study (GACRS) was a cross-sectional study of 1,165 children 6 to 14 years of age with asthma who were recruited between 2001 and 2011(6, 7). The study was approved by the Partners Human Research Committee at Brigham and Women’s Hospital (Boston, MA; protocol No. 2000-P-001130/55) and the Hospital Nacional de Niños (San José, Costa Rica).

Written informed consent was obtained from parents of all participants and assent was obtained from children from both studies. Whole blood DNA methylation profiling data was generated on 703 CAMP and 788 GACRS samples as part of the NHLBI Trans-Omics for Precision Medicine (TOPMed) program using the standard protocol for the Human Methylation 850K EPIC microarray BeadChips (Illumina, USA), as implemented at the University of Southern California Methylation Characterization Center.

**VDAART:**

The Vitamin D Antenatal Asthma Reduction Trial (VDAART) is a multi-center, randomized, and double-blinded study of the effect of vitamin D in pregnancy on risk of asthma in offspring. This study was conducted at three clinical centers across the United States between October 2009 and July 2011(8) and randomized 881 pregnant women to daily 4,400 IU (treatment arm) or 400 IU vitamin D (placebo or usual care arm) at 10 to 18 weeks’ gestation. Their offspring have been followed from delivery to childhood, now past age 6 years. Children had whole blood samples obtained at the ages of 1, 3, and 6 years and cord blood samples were obtained at delivery. Cord blood DNA methylation profiling data was generated on 572 children from the VDAART study for additional assessments at birth, using the 850K Illumina EPIC array at the Channing Division of Network Medicine, Brigham and Women’s Hospital, MA.

**Assessment of lung function levels**

Spirometry was done with an Eaglet Screener (ES) spirometer (Somerset Medical, Somerset, MA). Asthma was ascertained through verification of asthma history and medication use, as part of the inclusion criteria for GACRS and CAMP and subsequently verified with a methacholine challenge. In VDAART, the children were diagnosed with asthma/recurrent wheeze at age 3(9) and also a doctor’s asthma diagnosis at age 6(10) using parental reporting of a physician diagnosis which was based on the following five conditions: (1) wheeze after the child's second birthday, preceded by at least one report of wheeze prior to the second birthday; (2) use of asthma controller medication after the second birthday, preceded by wheeze before the second birthday; (3) two or more reports of wheeze after the second birthday; (4) at least one report of wheeze and use of asthma control medications at distinct visits after the second birthday; or (5) two distinct reports of use of asthma control medications after the second birthday. Lung function was measured in children by impulse oscillometer at 4 years of age and yearly after that and involved those who had completed at least two acceptable spirometry or impulse oscillometer tests and whose mother provided vitamin D measurements at baseline or during the third trimester. Spirometry was performed at the 5- and 6-year visits. Impulse oscillometer and spirometry were performed using the MasterScreen IOS and MasterScreen PFT systems with the Jaeger pneumotach (formerly CareFusion, now Vyaire Medical). Participants were instructed that the children should refrain from using bronchodilator inhalers, syrups, or pills for at least 8 hours before the visit and glucocorticoid inhalers or nebulizers and leukotriene modifiers for 24 hours before the visit. For spirometry, acceptability criteria were modified as preferred to allow for a minimum expiratory time of 1 second, with a 3-second expiratory time or a plateau in the volume–time curve.

***Statistical analyses***

**DNA-methylation data quality control and processing**

Data preprocessing and quality control was performed using minfi^38^. We used meffil^39^ for sample level outlier filtering. As part of the CpG site-based filtering, we removed probes with low detection pvalues (cut off 0.05), non-CpG probes (CH probes), probes containing SNPs at the CpG interrogation or at the single nucleotide extension and sex chromosomes. We calculated the blood cell count estimates using the Houseman algorithm^40,41^ using whole blood cell types as reference for GACRS and CAMP and cord blood cell types as reference for VDAART. Due to large number of sample batches, principal components from autosomes were specifically calculated from the DNA-methylation data. Background correction and data normalization was performed using functional normalization^42^ followed by type II probe and technical bias correction using Regression on Correlated probes (RCP) method^43^.

**Association of DNA-methylation with asthma and relevant LF phenotypes**

We tested CpGs against five LF phenotypes in each study: FEV_1_, FVC, FEV_1_/FVC ratio, FEF_25-75_ and FEF_25-75_/FVC ratio. In CAMP, two distinct longitudinal LF trajectories were analyzed: any ED (N=287) and any RG (N=339) compared to NG (N=183).

A multivariable robust linear regression model implemented in the robustbase R package was used to analyze single CpG site associations between DNA-methylation M-values as predictor and LF phenotypes in all three cohorts as outcome. A robust logistic regression model was used for the reduced growth and early decline LF trajectory phenotypes as outcome in CAMP and binary outcomes of asthma wheeze at age 3 and active asthma at age 6 in VDAART. All models were adjusted for age, sex, smoke exposure, height, body mass index, maternal asthma, methylation principal components associated with technical covariate sample plate and estimated cell type fractions. We additionally adjusted for self-reported race (African Americans, Whites, Hispanics, Others) provided by CAMP participants and child race/ethnicity and maternal gestational age in VDAART. In VDAART, if the LF data at age 6 was missing, then LF values at age 5 were used.

To understand sex-specificity and sex-divergent patterns, we further extended the EWAS models to evaluate the DNA-methylation associations with a sex interaction (DNA-methylation*sex) followed by a sex-stratified analysis. The differentially methylated CpGs from each study were examined for replication to identify early-life determinants of LF decline. An independent analysis of DNA-methylation signatures in sex chromosomes was performed for all LF phenotypes and in all populations to understand sex-based differences. To identify the most robust CpG marks, there is also utility in evaluating differentially methylated CpGs spanning a whole region (DMRs) to understand how global regional hypo- or hyper-methylation can regulate or maintain gene transcription. We used DMRCate^44^ to identify differentially methylated regions (DMRs) associated with LF phenotypes with defaults at an FDR threshold of 0.05.

IlluminaHumanMethylationEPICanno.ilm10b4.hg19 Bioconductor(11) package was used for the CpG to gene annotation with hg19/GRCh37 genome build for GACRS and CAMP and Infinium MethylationEPIC v1.0 B5 Manifest for VDAART.

**Gaussian Graphical Models (GGMs) on sex-stratified networks**

Our EWAS meta-analysis identified a set of 68 unique CpGs that were associated with at least one LF outcome and with opposing directions of effect in males and females in a stratified analysis. This analysis was different from the results where we identified differentially methylated positions (DMPs) that were significant only in males or females where we cannot compare the direction of effect. To investigate and target the relationships between these CpGs in males vs. in females, we constructed sex-stratified Gaussian graphical models (GGMs; also known as partial correlation networks) on the methylation of these CpGs in each of the three studies. Six total networks were thus estimated, among the following populations: males in GACRS, females in GACRS, males in CAMP, females in CAMP, males in VDAART, and females in VDAART. Methylation data as M values (logit base 2 of beta values) were used; subsequently, these data were transformed to approximate normality using the nonparanormal transformation as implemented in the npn function(12) from the R package huge(13). GGMs were estimated using the graphical lasso(14), also as implemented in huge. The optimal tuning parameter was selected using the extended Bayesian information criterion (eBIC), with hyperparameter gamma set to zero to encourage a denser model(15). Hub scores were calculated using the hub_score function(16) of the igraph package(17).

**Results**

**Epigenetic origins of lung function decline at birth and childhood**

Between the childhood cohorts GACRS and CAMP, 672 DMPs replicated for FEV_1_/FVC; 671 had consistent direction of effect; 587 were relatively hyper-methylated (87.5%) and 84 were relatively hypo-methylated (**Table E5)**. For FEF_25-75_, seven DMPs replicated, all with consistent effect; five were hyper-methylated and two were hypo-methylated (**Table E6**). For FEF_25-75_/FVC, 124 DMPs replicated, all with consistent effect; 100 were hyper-methylated (80.6%) and 24 were hypo-methylated (**Table E7)**. We did not identify any overlap for FEV_1_ and FVC.

Fewer associations replicated between VDAART, GACRS and CAMP (**Figure E2**). One CpG site cg07144755 annotated to *RGS3* gene was consistent in CAMP (Estimate=-0.52, FDR=3.8e-03) and VDAART (Estimate=-0.56, FDR=0.02) for the FEF_25-75_ LF outcome. One CpG site cg10730149 annotated to the *RPS6KC1* gene was consistent in GACRS (Estimate=-3.73, FDR=0.04) and VDAART (Estimate=-1.53, FDR=0.03) for the FEV_1_FVC LF outcome. There was no replication between all three cohorts at the CpG site level, however, at the gene-level, there was an overlap of 20, 2 (*RPTOR, PALLD*) and 4 (*RPTOR, C1orf21, PHACTR1, GPHN*) genes for FEV_1_/FVC, F_25-75_ and FEF_25-75_/FVC phenotypes respectively (**Figure E2**). Nine (*RUSC2, SEPT9, SGMS1, SIK3, SNX9, ST3GAL4, TRAPPC9, ZMIZ1*) of the 20 genes for FEV_1_FVC mapped to different CpGs but their methylation levels were consistent (hypo-methylated or hyper-methylated) in all three cohorts (**Table E8**). Four FEV_1_/FVC-associated DMPs (GACRS and CAMP, FDR<0.05) were also associated with asthma age 6y (VDAART, P<1e-03) mapping to cg23516680: *NOLC1, cg19752768: SWAP70, cg02359181: GPI, cg00005734: RALA* genes, implicated in allergic asthma and eosinophilia from the EWAS catalogue.

Encouraged by investigating temporal trends of LF growth and decline, we also explored the overlap between lung function trajectory DMPs identified in CAMP and DMPs associated with asthma outcomes in VDAART to identify potential targets for asthma development (**Figure E2F**). We found 338 and 534 significant DMPs associated with any early LF decline and reduced growth compared to normal growth trajectories in CAMP (1x10^-3^) (**Table 2**). We identified one DMP (*ALPK2* gene) negatively associated with any early decline (Estimate=-1.79, p-value=8.6e-04), reduced growth (Estimate=-1.68, p-value=6.0e-04), and active asthma outcome (Estimate=-2.59, p-value=9.8e-04). Two CpGs: cg12728791 annotated to *HMCN1* gene and cg09234670 annotated to *NQO2* gene (**Figure E3**) were shared between reduced growth and asthma outcome (VDAART: Estimate=-2.75, p-value=8.8e-04; CAMP_RG_: Estimate=-1.1, p-value=4.8e-04) and early decline and asthma outcome (VDAART: Estimate=2.54, p-value=3.5e-04; CAMP_ED_: Estimate=0.99, p-value=4.7e-04) respectively.

We further identified several LF-associated DMPs for the sex-specific overlap and the between sex differences during adolescence than at birth. From the interaction model (DNAm*sex), we identified DMPs in CAMP and VDAART, but not in GACRS (**Table E9**). Stratifying by sex identified several LF-DMPs specific to males or females (**Table E9**). Only one CpG site cg07762234 (C16orf79 gene) associated with FEV_1_ in CAMP overlapped between the interaction model (Estimate_males_=-0.49, FDR=4.27e-06) and the sex-stratified models in both males (Estimate=0.34, FDR=3.1e-04) and females (Estimate=0.31, FDR=2.94e-05) with divergent DNA-methylation levels. Only few consistent DMPs replicated between cohorts and in opposite direction of effect between males and females within each cohort (FDR<0.05, **Table E10**). We identified two male-specific (FEV_1_, cg14294664:*ZNF644;* FEF_25-75_/FVC, cg11297046:*HRC*) and two female-specific CpGs (FEV_1_/FVC, cg23067272:*U2;* FEF_25-75_, cg01105494:*SPAG17*) that replicated and in consistent direction of methylation effect between VDAART and GACRS (**Table E10**). We identified one male-specific (FEV_1_, cg02298956:*ZNF772*) and three female-specific CpGs (FEV_1_/FVC, cg08158518:*SPSB4* and cg09641127:*KLF11*; FEF_25-75_/FVC, cg14094882: *TRIP4*) that replicated and in consistent direction of methylation effect between VDAART and CAMP (**Table E10**). Between cohorts, we identified a few sex-divergent patterns and only during childhood (**Table E10**) in CRA (FVC, cg26436330:*COL11A1*; FEF_25-75_/FVC, cg04883139:*NCK1*) and CAMP (FEF_25-75_/FVC, cg21742872:*C2CD2*; cg00857557:*MEA1*; cg08071719:*MCF2L*).

From the regional analysis, we identified 5, 262, 142 and 250 significantly associated DMRs for FEV_1_, FEV_1_/FVC, FEF_25-75_ and FEF_25-75_/FVC respectively in CAMP at a Stouffer’s FDR<0.05. We identified 153 and 19 significantly associated DMRs for FEV_1_/FVC and FEF_25-75_/FVC at a Stouffer’s FDR<0.05 in GACRS. The overlap of these DMRs between GACRS and CAMP is included in the main manuscript highlighting common replicated childhood specific DMRs.

***Epigenetic age acceleration and LF decline***

Epigenetic age acceleration was significantly associated with reduction in all LF outcomes in CAMP. This trend stayed robust when stratified by sex, mostly for females where the reduction was more evident compared to males (**Table E18**). There was a suggestive association for age acceleration in COPD subjects in the reduced growth trajectory (Wilcoxon p-value=0.09), which did not stay significant when adjusted for covariates. When stratified by sex, the trend stayed suggestively significant in females in the adjusted model (coefficient=0.15, p-value=0.078) and was significant in females (coefficient=0.23, p-value=0.03) considered on a reduced growth/early decline trajectory compared to those on normal trajectory. This latter effect was significant after adjusting for the epigenetic age acceleration*maternal asthma interaction term suggesting conditional dependence on maternal asthma. Further, a one-unit increase in age acceleration residual was associated with a higher odd (relative risk ratio) of being in the mild COPD category in males and moderate COPD in females compared to no COPD, however this association remained suggestive after adjusting for all covariates (**Table E18**). In more homogeneous GACRS, the trend was not significant. In cord blood from the VDAART study, epigenetic age acceleration was significantly associated with an increased FEV_1_ and FVC by age 5-6.

**Supplementary Tables**

**Table E1.** Characteristics of the study participants in the CAMP cohort with data on DNA methylation, lung function outcomes and trajectories of growth and decline

| Characteristics | Total Samples (N=703) | NG  (N=183) | Any ED  (N=287) | Any RG  (N=339) | P-Value (NG-ED)^*^ | P-Value (NG-RG)^*^ |
| --- | --- | --- | --- | --- | --- | --- |
| Age_F48_, mean (SD) | 12.9 (2.1) | 12.6 (2.2) | 13.2 (2.0) | 12.6 (2.1) | 3.1x10^-3^ | 0.14 |
| Sex, n (%) |  |  |  |  | 0.24 | 1.2x10^-3^ |
| Males | 423 (60.2) | 97 (53) | 169 (58.9) | 228 (67.3) |  |  |
| Females | 280 (39.8) | 86 (47) | 118 (41.1) | 111 (32.7) |  |  |
| ETS, n (%) |  |  |  |  | 0.48 | 0.58 |
| Absence | 424 (60.3) | 109 (59.6) | 163 (56.8) | 215 (63.4) |  |  |
| Presence | 276 (39.3) | 71 (38.8) | 124 (43.2) | 124 (36.6) |  |  |
| BDR %, mean (SD) | 0.10 (0.09) | 0.07 (0.06) | 0.09 (0.08) | 0.12 (0.10) | 0.02 | 9.5x10^-7^ |
| PRE BD FEV_1_, mean (SD) | 2.6 (0.8) | 2.7 (0.7) | 2.7 (0.8) | 2.4 (0.7) | 0.31 | 6.5x10^-6^ |
| POST BD FEV_1_, mean (SD) | 2.8 (0.8) | 2.9 (0.8) | 3.0 (0.8) | 2.6 (0.8) | 0.17 | 4.9x10^-4^ |
| PRE BD FVC, mean (SD) | 3.3 (1.0) | 3.31 (0.95) | 3.50 (1.01) | 3.18 (0.96) | 0.04 | 0.14 |
| POST BD FVC, mean (SD) | 3.4 (1.0) | 3.34 (0.96) | 3.55 (1.02) | 3.23 (0.96) | 0.02 | 0.25 |
| PRE BD FEV_1_/FVC, mean (SD) | 77.9 (8.9) | 81.1 (7.0) | 78.7 (8.8) | 74.8 (9.2) | 9.3x10^-4^ | <2.2x10^-16^ |
| POST BD FEV_1_/FVC, mean (SD) | 83.7 (7.0) | 86.0 (5.6) | 84.0 (6.9) | 81.3 (7.4) | 1.9x10^-4^ | 7.4x10^-16^ |
| PRE BD FEF_25-75_, mean (SD) | 2.34 (0.95) | 2.57 (0.85) | 2.55 (1.02) | 1.97 (0.82) | 0.78 | 1.2x10^-13^ |
| POS BD FEF_25-75_, mean (SD) | 2.9 (1.0) | 3.2 (0.9) | 3.1 (1.0) | 2.5 (0.9) | 0.65 | 1.4x10^-12^ |
| PRE BD FEF_25-75_/FVC, mean (SD) | 0.7 (0.2) | 0.8 (0.2) | 0.7 (0.2) | 0.6 (0.2) | 9.3x10^-3^ | 1.4x10^-14^ |
| POST BD FEF_2575_/FVC, mean (SD) | 0.9 (0.2) | 0.97 (0.2) | 0.90 (0.2) | 0.8 (0.2) | 6.0x10^-4^ | 1.0x10^-14^ |
| BMI, mean (SD) | 21.5 (4.7) | 21.2 (4.2) | 23.0 (4.9) | 20.4 (4.3) | 2.7x10^-3^ | 0.16 |
| Race, n (%) |  |  |  |  | 6.4x10^-4^ | 3.4x10^-4^ |
| White | 486 (69.1) | 142 (77.6) | 182 (63.4) | 223 (65.8) |  |  |
| African Americans | 90 (12.8) | 26 (14.2) | 40 (13.9) | 37 (10.9) |  |  |
| Hispanic | 66 (9.4) | 8 (4.4) | 40 (13.9) | 44 (13.0) |  |  |
| Other | 61 (8.7) | 7 (3.8) | 25 (8.7) | 35 (10.3) |  |  |
| Treatment, n (%) |  |  |  |  | 0.22 | 0.93 |
| Placebo | 285 (40.5) | 79 (43.2) | 112 (39.0) | 135 (39.8) |  |  |
| Nedocramil | 207 (29.4) | 45 (24.6) | 92 (32.1) | 101 (29.8) |  |  |
| Budenoside | 211 (30.0) | 59 (32.2) | 83 (28.9) | 103 (30.4) |  |  |
| Height_cm_, mean (SD) | 156 (13.5) | 154 (12.8) | 158.5 (13.6) | 153.7 (13.9) | 3.2x10^-4^ | 0.16 |
| ICS use, n (%) |  |  |  |  | 0.81 | 0.65 |
| No | 432 (61.5) | 117 (63.9) | 180 (62.7) | 209 (61.7) |  |  |
| Yes | 267 (38.0) | 65 (35.5) | 107 (37.3) | 129 (38.1) |  |  |
| Maternal asthma, n (%) |  |  |  |  | 0.04 | 0.03 |
| No | 510 (72.5) | 156 (85.2) | 113 (39.4) | 139 (41.0) |  |  |
| Yes | 177 (25.2) | 34 (18.6) | 44 (15.3) | 62 (18.3) |  |  |

Abbreviations: BDR: bronchodilator response; ETS: environmental tobacco smoke; ICS: inhaled corticosteroid use; BMI: body mass index; SD: standard deviation; CAMP: Childhood Asthma Management Program; NG: normal growth; ED: early decline; RG: reduced growth; Missingness: Data for 13 subjects was missing for age at F48; data for 3 subjects was missing for ETS; data for 19 subjects was missing for BD change % at F48; data for 18 subjects was missing for BMI at F48; data for 16 subjects was missing for height at F48; data for 16 subjects was missing for maternal asthma. N varies for lung function phenotypes

* Significance of difference was evaluated using chi-squared test for categorical variables and two-sample t-test for continuous variables.

**Table E2.** Characteristics of the GACRS study participants with DNA methylation data and data on lung function

| Characteristics | Total Samples (N=788) |
| --- | --- |
| Age, mean (SD) | 9.3 (1.9) |
| Sex, n (%) |  |
| Males | 467 (59.3) |
| Females | 321 (40.7) |
| ETS, n (%) |  |
| Absence | 543 (68.9) |
| Presence | 240 (30.5) |
| BDR %, mean (SD) | 5.7 (10.4) |
| PRE BD FEV_1_, mean (SD) | 1.8 (0.5) |
| POST BD FEV_1_, mean (SD) | 1.9 (0.5) |
| PRE BD FVC, mean (SD) | 2.1 (0.6) |
| POST BD FVC, mean (SD) | 2.2 (0.6) |
| PRE BD FEV_1_/FVC, mean (SD) | 83.3 (7.5) |
| POST BD FEV_1_/FVC, mean (SD) | 86.3 (6.7) |
| PRE BD FEF_25-75_, mean (SD) | 2.0 (0.7) |
| POS BD FEF_25-75_, mean (SD) | 2.3 (0.8) |
| PRE BD FEF_25-75_/FVC, mean (SD) | 0.9 (0.3) |
| POST BD FEF_2575_/FVC, mean (SD) | 1.1 (0.3) |
| BMI, mean (SD) | 18.4 (3.8) |
| Race, n (%) |  |
| White | 788 (100) |
| Height_cm_, mean (SD) | 133 (11.7) |
| Maternal asthma, n (%) |  |
| No | 545 (69.2) |
| Yes | 240 (30.5) |

Abbreviations: BDR: bronchodilator response; ETS: environmental tobacco smoke; ICS: inhaled corticosteroid use; BMI: body mass index; SD: standard deviation; CRA: The Genetic Epidemiology of Asthma in Costa Rica Study; Missingness: Data for three subjects was missing for BMI and height; data for three subjects was missing for maternal asthma; data for five subjects was missing for smoke exposure; N varies for lung function phenotypes

**Table E3.** Characteristics of the VDAART study participants with DNA methylation data and data on lung function and asthma outcomes

| Characteristics | Total Samples (N=572) |
| --- | --- |
| Gestational age in weeks, mean (SD) | 38.8 (1.5) |
| Gestational age, n (%) |  |
| <37 weeks | 36 (6.3) |
| >=37 weeks | 536 (93.7) |
| Child sex, n (%) |  |
| Males | 304 (53.1) |
| Females | 268 (46.9) |
| Smoke exposure, n (%) |  |
| Absence | 559 (97.7) |
| Presence | 13 (2.3) |
| Best/PRE FEV_1_, mean (SD) | 1.2 (0.2) |
| Best/PRE FVC, mean (SD) | 1.4 (0.3) |
| Best/PRE FEV_1_/FVC, mean (SD) | 0.9 (0.1) |
| Best/PRE BD FEF_25-75_, mean (SD) | 1.6 (0.4) |
| Best/PRE FEF_25-75_/FVC, mean (SD) | 1.2 (0.3) |
| Asthma wheeze at age 3, n (%) |  |
| Absence | 387 (67.7) |
| Presence | 145 (25.3) |
| Asthma at age 6, n (%) |  |
| Absence | 424 (74.1) |
| Presence | 84 (14.7) |
| Race, n (%) |  |
| White | 202 (35.3) |
| African Americans | 253 (44.2) |
| Others | 117 (20.5) |
| Height_m_, mean (SD) | 1.2 (0.1) |
| Maternal Vitamin D in 3^rd^ trimester, mean (SD) | 33.5 (14.4) |

Abbreviations: SD: standard deviation; VDAART: The Vitamin D Antenatal Asthma Reduction Trial; Missingness: Data for 252 subjects was missing for FEV1, FVC, FEV1FVC, FEF25-75 and FEF25-75/FVC lung function phenotypes; data for 40 subjects was missing for asthma wheeze at age 3; data for 64 subjects was missing for asthma at age 6; data for maternal vitamin D in third trimester (week 32-38) was missing for nine subjects; N varies for lung function phenotypes

**Table E4.** Number of epigenome-wide LF-associated differentially methylated positions (DMPs) individually in all three studies GACRS, CAMP and VDAART and replication between them

**Table E4A.** DMPs in relation to asthma, pre-bronchodilator lung function and trajectory outcomes at different statistical thresholds in all three studies

| Cohorts and LF/asthma phenotype outcomes | Associations at 1e-03 | Associations at FDR | Genome-wide threshold |
| --- | --- | --- | --- |
| CAMP |  |  |  |
| LLF outcome |  |  |  |
| NG vs. Any RG | 534 | 0 | 0 |
| NG vs. Any ED | 338 | 0 | 0 |
| LF outcomes |  |  |  |
| FEV_1_ | 3,229 | 387 | 21 |
| FVC | 1,743 | 89 | 19 |
| FEV1/FVC | 6,099 | 3,485 | 328 |
| FEF2575 | 6,181 | 3,463 | 293 |
| FEF2575/FVC | 6,613 | 4,033 | 427 |
| GACRS |  |  |  |
| FEV_1_ | 1,490 | 38 | 12 |
| FVC | 1,815 | 45 | 11 |
| FEV_1_/FVC | 9,910 | 3,699 | 38 |
| FEF_25-75_ | 4,261 | 160 | 14 |
| FEF_25-75_/FVC | 6,480 | 801 | 29 |
| VDAART |  |  |  |
| Last FEV_1_ | 2,218 | 359 | 78 |
| Last FVC | 6,681 | 1,043 | 58 |
| Last FEV_1_/FVC | 2,842 | 233 | 39 |
| Last FEF_25-75_ | 2,439 | 346 | 56 |
| Last FEF_25-75_/FVC | 3,003 | 414 | 62 |
| Asthma/wheeze at age 3 | 587 | 0 | 0 |
| Asthma at age 6 | 5,936 | 0 | 0 |

Abbreviations: LF, lung function; LLF, longitudinal trajectory outcomes in CAMP; NG, normal growth; RG, reduced growth; ED, early decline; FDR, false discovery rate

**Table E4B.** Number of LF-associated DMPs by sex (FDR<0.05).

| **Cohorts and significant LF associations** | **Associations (with sex interaction)** | **Associations at FDR (sex-stratified)** | | **Overlap of associations^+^** | |
| --- | --- | --- | --- | --- | --- |
| **CAMP** | **CpG*Sex** | **M** | **F** | **M** | **F** |
| FEV_1_ | 37 | 357 | 723 | 7 | 14 |
| FVC | 73 | 212 | 469 | 2 | 6 |
| FEV_1_/FVC | 0 | 565 | 1997 | 0 | 0 |
| FEF_25-75_ | 10 | 776 | 2121 | 5 | 4 |
| FEF_25-75_/FVC | 1 | 1151 | 2028 | 0 | 1 |
| **GACRS** | **CpG*Sex** | **M** | **F** | **M** | **F** |
| FEV_1_ | 0 | 261 | 253 | 0 | 0 |
| FVC | 0 | 261 | 276 | 0 | 0 |
| FEV_1_/FVC | 0 | 909 | 401 | 0 | 0 |
| FEF_25-75_ | 0 | 236 | 163 | 0 | 0 |
| FEF_25-75_/FVC | 0 | 371 | 869 | 0 | 0 |
| **VDAART** | **CpG*Sex** | **M** | **F** | **M** | **F** |
| Last FEV_1_ | 66 | 364 | 702 | 21 | 6 |
| Last FVC | 122 | 518 | 574 | 21 | 8 |
| Last FEV_1_/FVC | 16 | 813 | 598 | 6 | 5 |
| Last FEF_25-75_ | 11 | 551 | 1091 | 5 | 3 |
| Last FEF_25-75_/FVC | 4 | 864 | 1393 | 2 | 1 |

Models were adjusted for age, smoke exposure, height, body mass index, maternal asthma, principal components and cell type fractions and additionally for race in CAMP and child race in VDAART

**^+^**The number of overlapping associations between the interaction model and the sex-stratified models for males and females.

**Table E4C**. Sex-specific lung function associations between (DNA-methylation in cord blood and during childhood) and within cohorts. The CpGs were reported here if they were in the same direction between cohorts and in opposite direction of effect between sexes within cohorts.

| CpG | Lung Function | CHR | CpG position | Gene | Estimate | FDR P-value | Function and Implications |
| --- | --- | --- | --- | --- | --- | --- | --- |
| *Consistent within phenotype associations across populations (VDAART and GACRS)* | | | | | | |  |
| cg14294664-M | FEV_1_ | 1 | 91,448,818 | *ZNF644* | 0.08, 0.05 | 5.6e-04, 0.02 | Transcriptional regulation, Increased expression in Neutrophilic asthma, genetics of asthma progression |
| cg23067272-F | FEV_1_/FVC | 11 | 111,252,327 | *U2* | -1.75, -2.50 | 0.05, 8.9e-03 | Small nuclear RNA, inflammation and RNA splicing |
| cg01105494-F | FEF_25-75_ | 1 | 118,727,908 | *SPAG17* | -0.25, -0.21 | 0.03, 6.5e-03 | Organization of motile cilia in respiratory tract, Hydroxy-methylation and deregulated in allergic asthma mice (early life and adulthood) and neutrophilic asthma |
| cg11297046-M | FEF_25-75_/FVC | 19 | 49,657,000 | *HRC* | -0.07, -0.28 | 0.05, 0.05 | Insulin Growth Factor regulation, calcium ion binding, Tumor metastasis |
| *Consistent within phenotype associations across populations (VDAART and CAMP)* | | | | | | |  |
| cg02298956-M | FEV_1_ | 19 | 57,989,134 | *ZNF772* | -0.12, -0.29 | 0.04, 0.03 | Regulation of transcription factor activity, genetic studies of asthma |
| cg08158518-F | FEV_1_/FVC | 3 | 140,866,589 | *SPSB4* | 1.08, 2.19 | 8.2e-08, 0.03 | protein polyubiquitination and Class I MHC-mediated antigen processing |
| cg09641127-F |  | 2 | 10,193,895 | *KLF11* | -6.05, -6.04 | 0.03, 0.04 | Transcription factor, mediator of TGF-beta, lung cancer, DNA methylation associated with asthma severity |
| cg14094882-F | FEF_25-75_/FVC | 15 | 64,680,186 | *TRIP4* | -0.11, -1.88 | 0.05, 0.03 | Transcriptional coactivator with *EP300, CREBBP and NCOA1*; Thyroid and estrogen receptor binding, high in non-small cell lung cancer |
| *Sex-specific phenotype associations within populations (males and females)* | | | | | | |  |
| GACRS |  |  |  |  |  |  |  |
| cg26436330 | FVC | 1 | 103,573,700 | *COL11A1* | -0.08, 0.09 | 8.7e-03, 7.5e-05 | Extracellular matrix binding, promotes non-small cell lung cancer |
| cg04883139 | FEF_25-75_/FVC | 3 | 136580967 | *NCK1* | -0.11, 0.13 | 0.04, 7.3e-04 | Cell adhesion, cytoskeletal dynamics, progression of lung cancers |
| CAMP |  |  |  |  |  |  |  |
| cg21742872 | FEF_25-75_/FVC | 21 | 43,373,869 | *C2CD2* | 0.12, -0.09 | 0.02, 0.01 | Calcium and phosphoinositide signaling, skeletal disorders |
| cg00857557 |  | 6 | 42,982,050 | *MEA1* | 0.11, -0.14 | 0.04, 9.28e-05 | Testicular differentiation, Covid19 susceptibility in lung cancer |
| cg08071719 |  | 13 | 113,650,155 | *MCF2L* | -0.30, 0.09 | 0.04, 0.02 | guanyl-nucleotide exchange factor activity and 1-phosphatidylinositol binding, cardiovascular, bone and brain disorders, lung cancer |

**Tables E5-E10 and E12-S18 in excel file due to size.**

**Table E11.** Measures of epigenetic age acceleration and significant associations with lung function outcomes

| ***Epigenetic clock metrics in GACRS, CAMP (F48) and VDAART*** | | |
| --- | --- | --- |
|  | **Age Acceleration residual** | |
| **Multivariate model outcomes (EN Clock)** | | |
| **CAMP** | **Estimate (95% CI)** | **P-value** |
| PRFEV_1_ | -0.02 (-0.03, -0.01) | **3.3x10^-3^** |
| Males | -0.020 (-0.04, -0.01) | **0.03** |
| Females | -0.019 (-0.04, -0.00) | **0.05** |
| PREFEV_1_/FVC | -0.37 (-0.65, -0.09) | **0.01** |
| Males | -0.16 (-0.54, 0.21) | 0.39 |
| Females | -0.73 (-1.17, -0.29) | **0.001** |
| PRE_25-75_ | -0.03 (-0.06, -0.01) | **0.02** |
| Males | -0.02 (-0.06, 0.01) | 0.23 |
| Females | -0.05 (-0.09, -0.01) | **6.4x10^-3^** |
| PRE_25-75_/FVC | -0.00007 (-0.00, -0.01) | **0.04** |
| Males | -0.003 (-0.01, 0.01) | 0.59 |
| Females | -0.02 (-0.03, -0.01) | **5.1x10^-3^** |
| COPD Gold stages (males_0,1_) | RR ratio: 0.58 | 0.08 |
| COPD Gold stages (females_0,2_) | RR ratio: 0.46 | 0.06 |
| **GACRS** | **Estimate (95% CI)** | **P-value** |
| PRFEV_1_ | -0.003 (-0.01, 0.01) | 0.30 |
| PREFEV_1_/FVC | -0.08 (-0.24, 0.07) | 0.29 |
| PRE_25-75_ | -0.01 (-0.02, 0.01) | 0.30 |
| PRE_25-75_/FVC | -0.002 (-0.01, 0.01) | 0.50 |
| **VDAART (EPIC clock)** | **Estimate (95% CI)** | **P-value** |
| FEV_1_ | 0.03 (0.005, 0.06) | **0.04** |
| FVC | 0.03 (0.005, 0.06) | **0.03** |

Abbreviations and notes: All models were adjusted for the same covariates as included for the EWAS models.

CI: confidence interval; EN clock: elastic net clock; In CAMP, GOLD stages were defined as follows: GOLD stage 1 - mild: FEV1 ≥80% predicted. GOLD stage 2 - moderate: 50% ≤ FEV1 <80% predicted. GOLD stage 3 - severe: 30% ≤ FEV1 <50% predicted. GOLD stage 4 - very severe: FEV1 <30% predicted. RR ratio: The relative risk ratio/log odds for a one-unit increase in AgeAccelerationResidual for being in mild, moderate, or severe COPD category compared to normal.

**Supplementary Figures**

**Figure E1.** Venn diagram overlap of DNA-methylation associations with all lung function phenotypes in all three studies. A) GACRS, B) CAMP and C) VDAART

A) GACRS

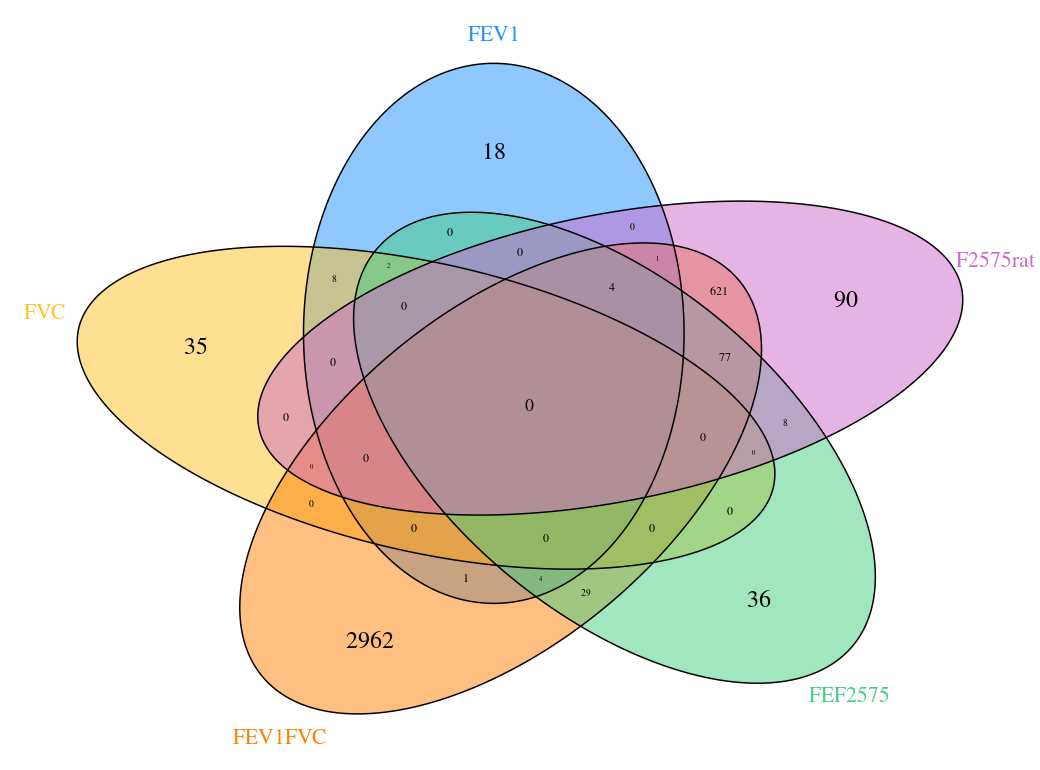

B) CAMP

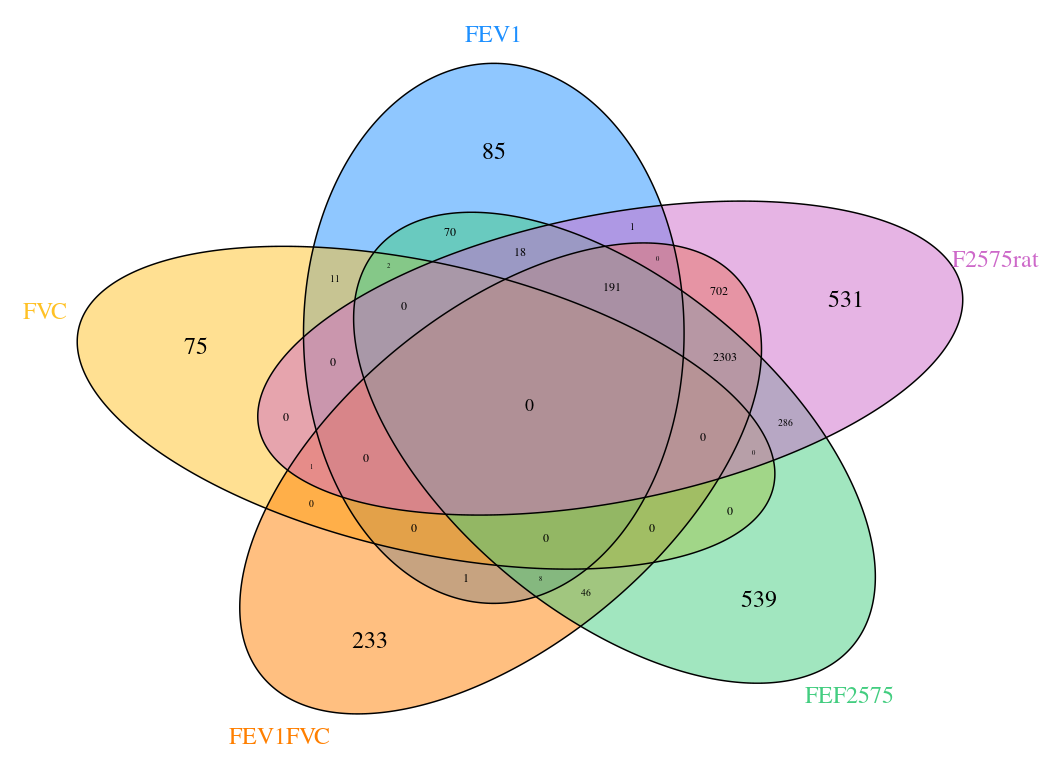

C) VDAART

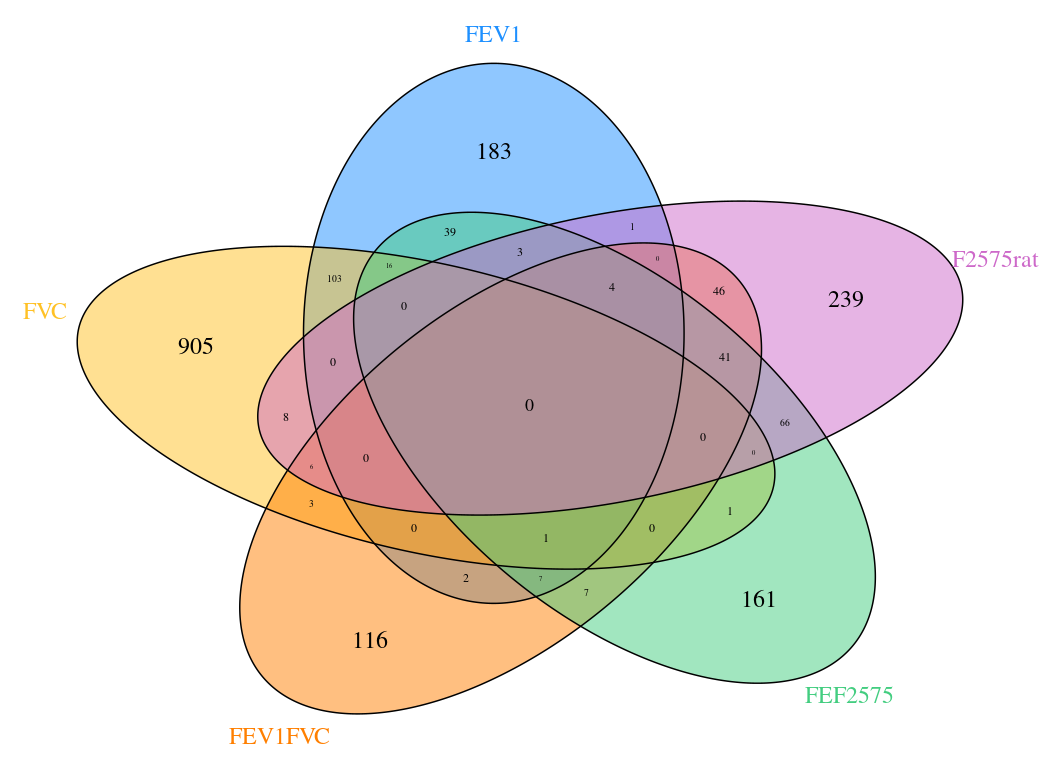

**Figure E2.** Intersection of lung function associated CpGs and their annotated genes from individual study-level EWAS across all cohorts (CRA same as GACRS study).

A) Upset Plots for the lung function phenotype FEV_1_ showing intersecting/overlapping CpGs (left) and their annotated genes (right) between all three cohorts.

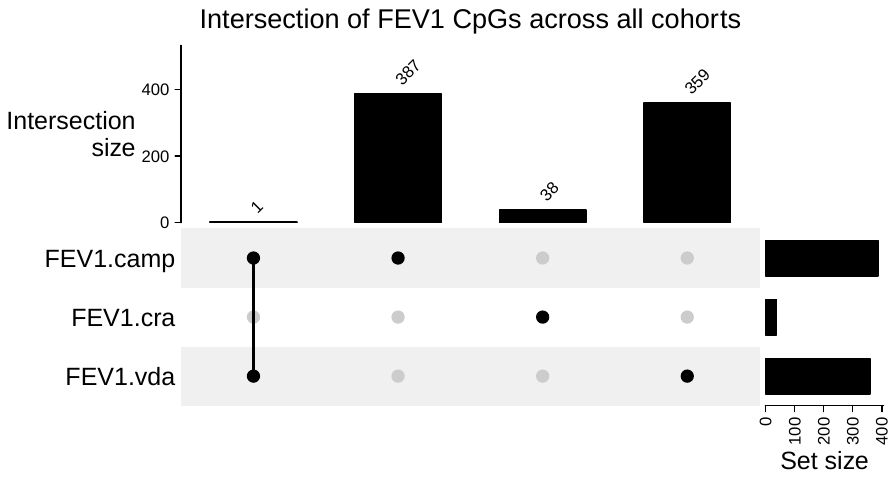

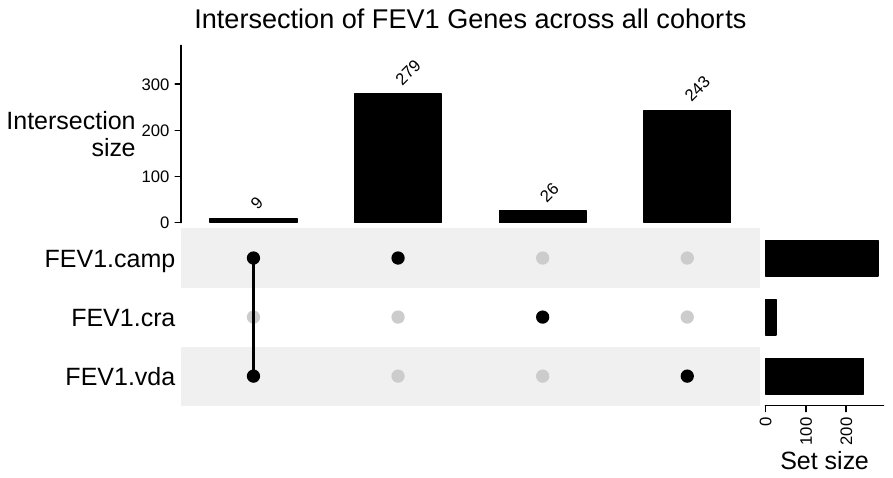

B) Upset Plots for the lung function phenotype FVC showing intersecting/overlapping CpGs (left) and their annotated genes (right) between all three cohorts.

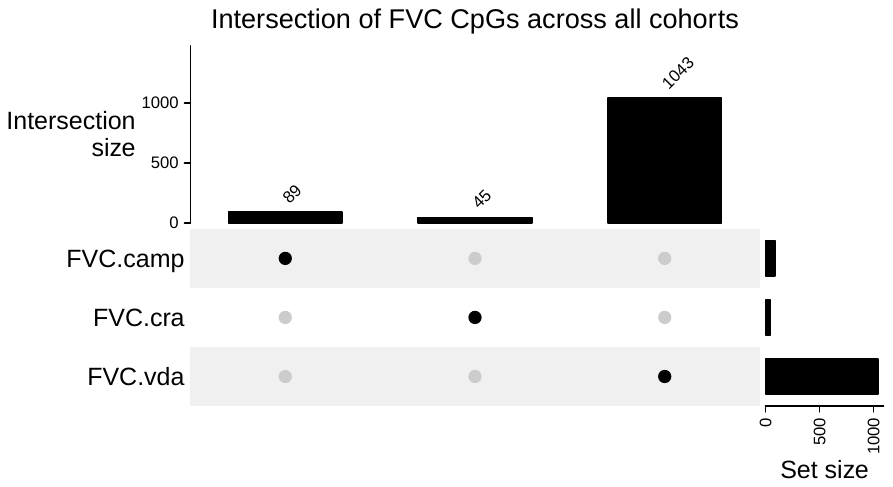

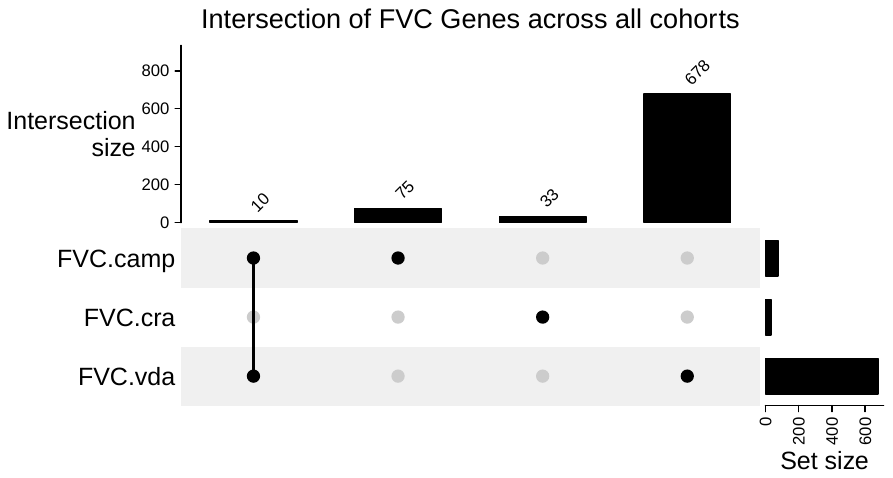

C) Upset Plots for the lung function phenotype FEV_1_/FVC showing intersecting/ overlapping CpGs (left) and their annotated genes (right) between all three cohorts.

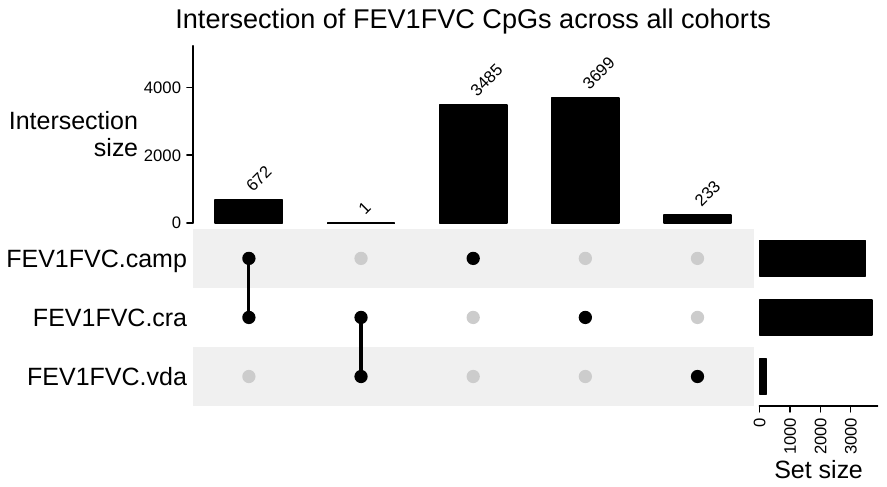

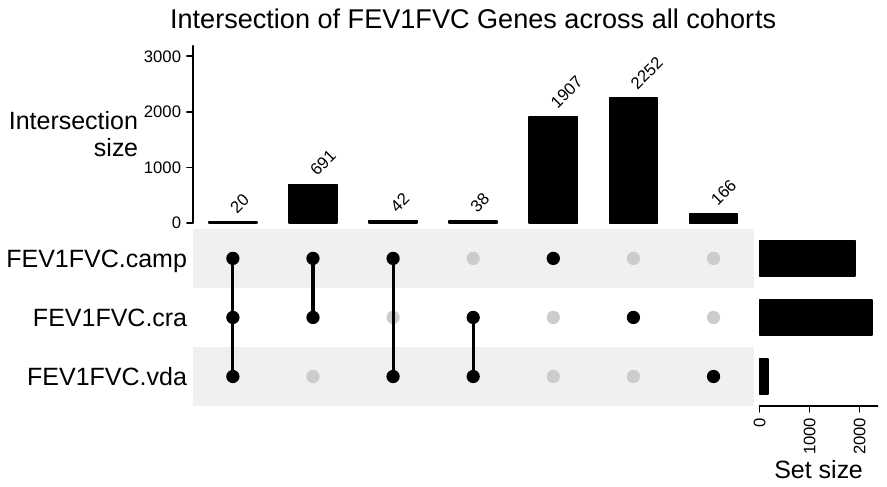

D) Upset Plots for the lung function phenotype FEF_25-75_ showing intersecting/ overlapping CpGs (left) and their annotated genes (right) between all three cohorts.

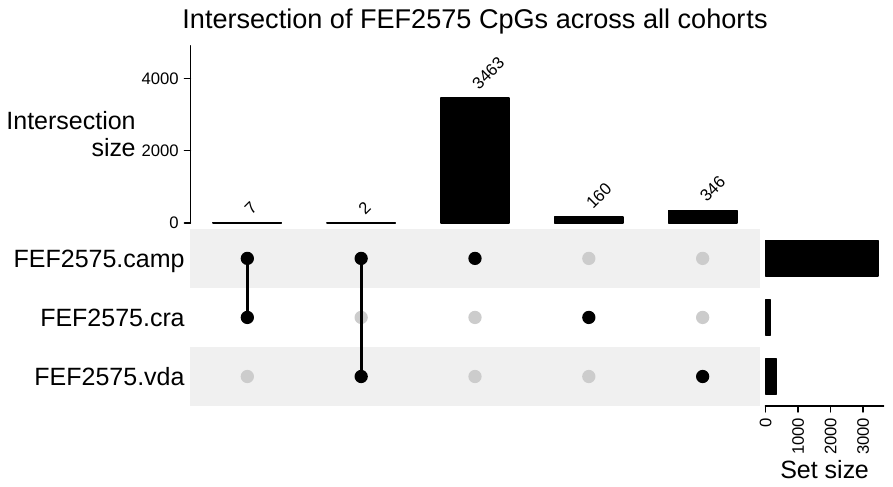

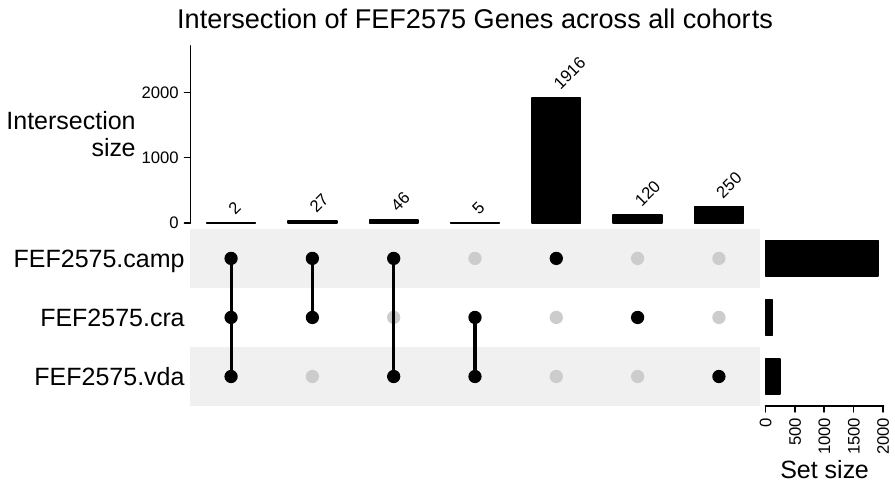

E) Upset Plots for the lung function phenotype FEF_25-75_/FVC showing intersecting/ overlapping CpGs (left) and their annotated genes (right) between all three cohorts.

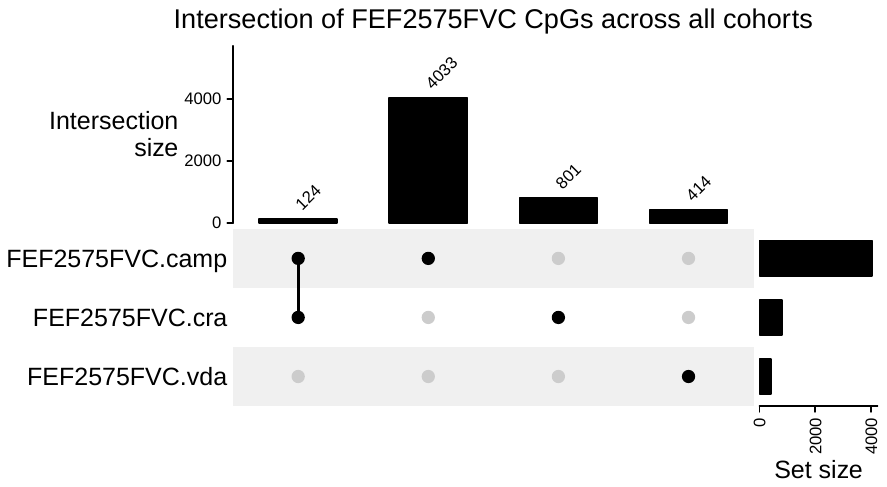

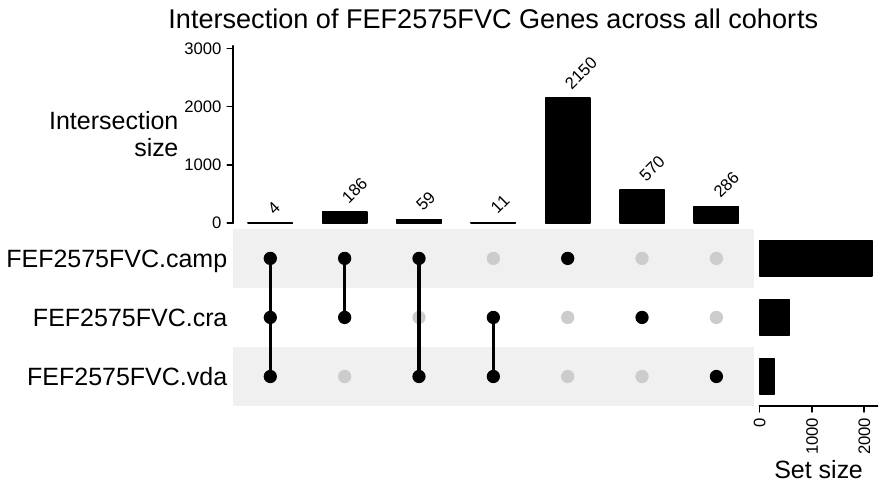

F) Upset Plots for the lung function trajectories in CAMP and asthma outcomes in VDAART showing the intersection of CpGs (left) and their annotated genes (right).

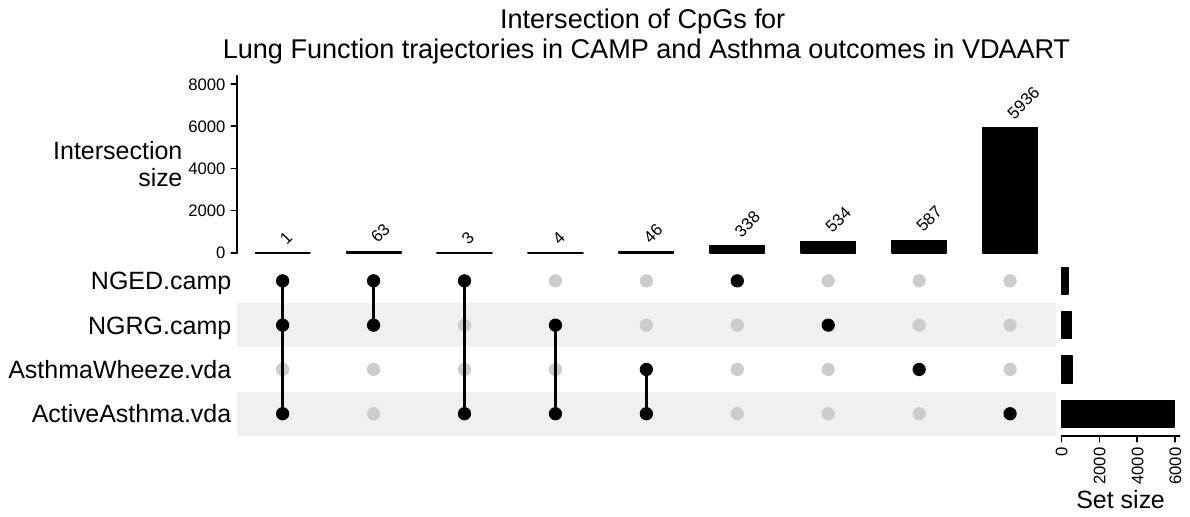

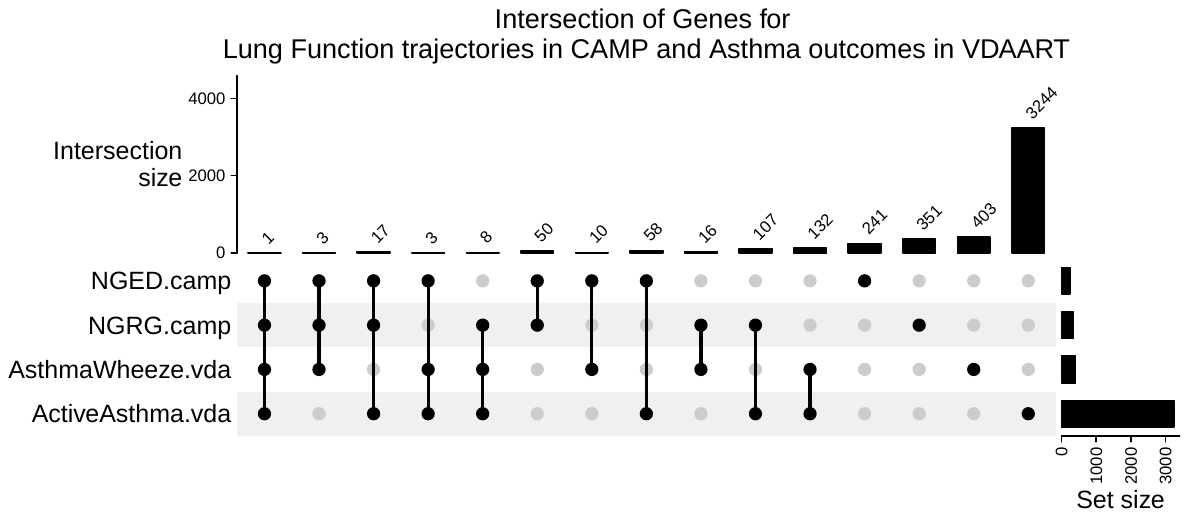

**Figure E3.** Overlap between lung function trajectories in CAMP and asthma outcomes in VDAART

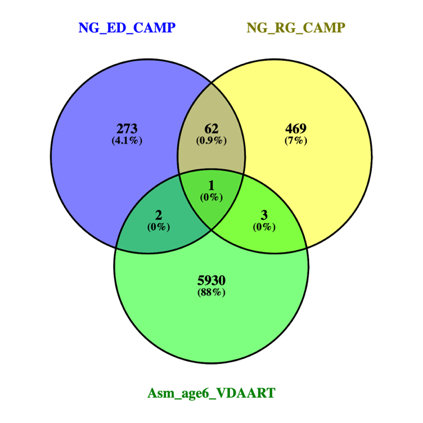

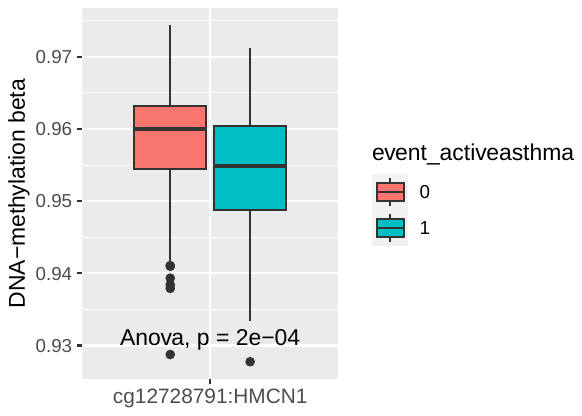

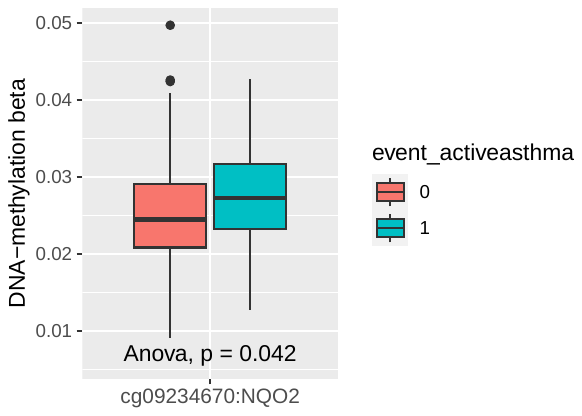

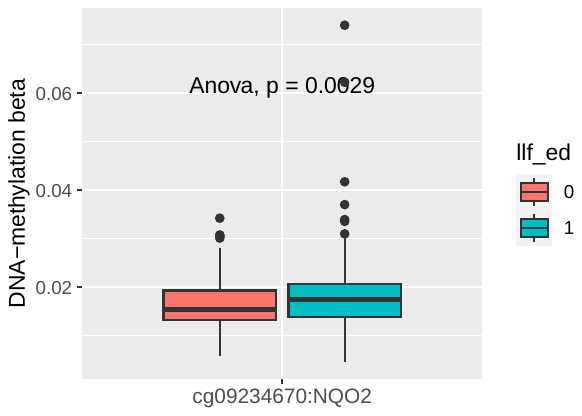

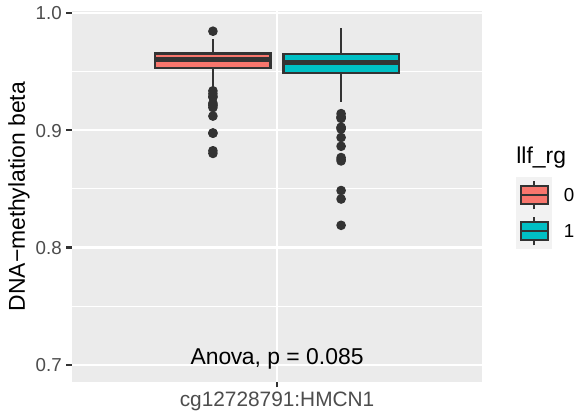

cg09234670: NQO2

cg19257719: ALPK2

cg12728791: HMCN1

cg19257719: ALPK2

**Figure E4.** Overlap of the lung function associations after adjustment of EWAS associations and effect modification by polygenic risk scores. The Upset plots show the intersect mode highlighting the significant associations from the original EWAS models, followed by the models that adjusted for PRS and the models that included the interaction term (CpG * PRS) for FEV_1_ and FEV_1_/FVC.

A. **B.**

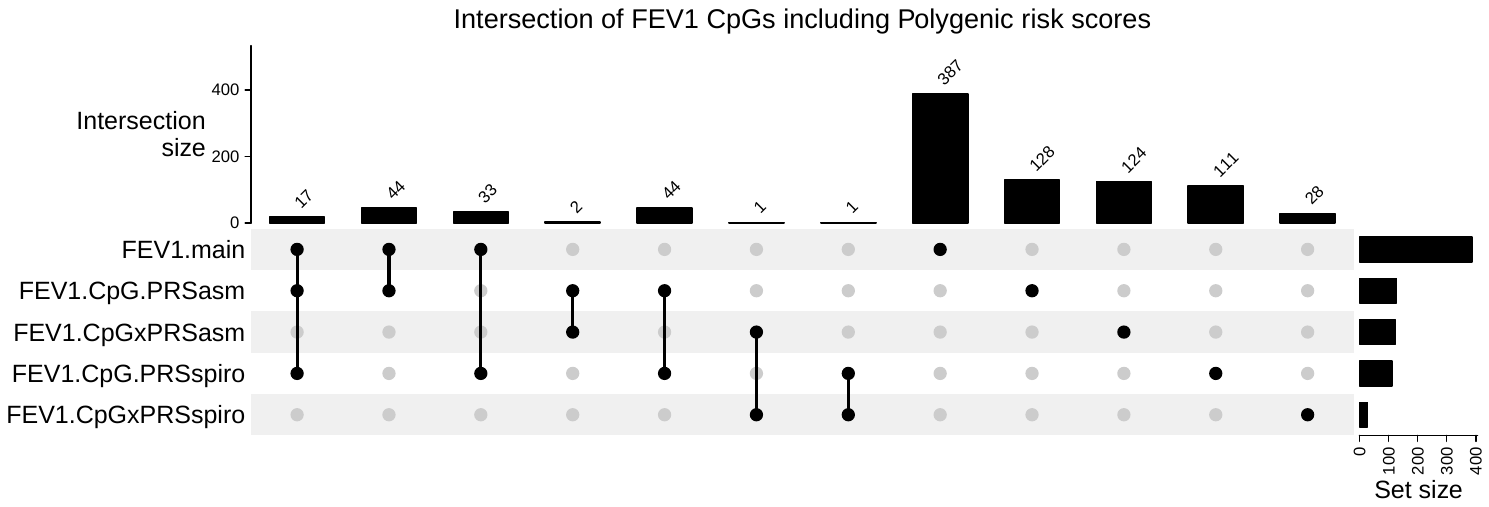

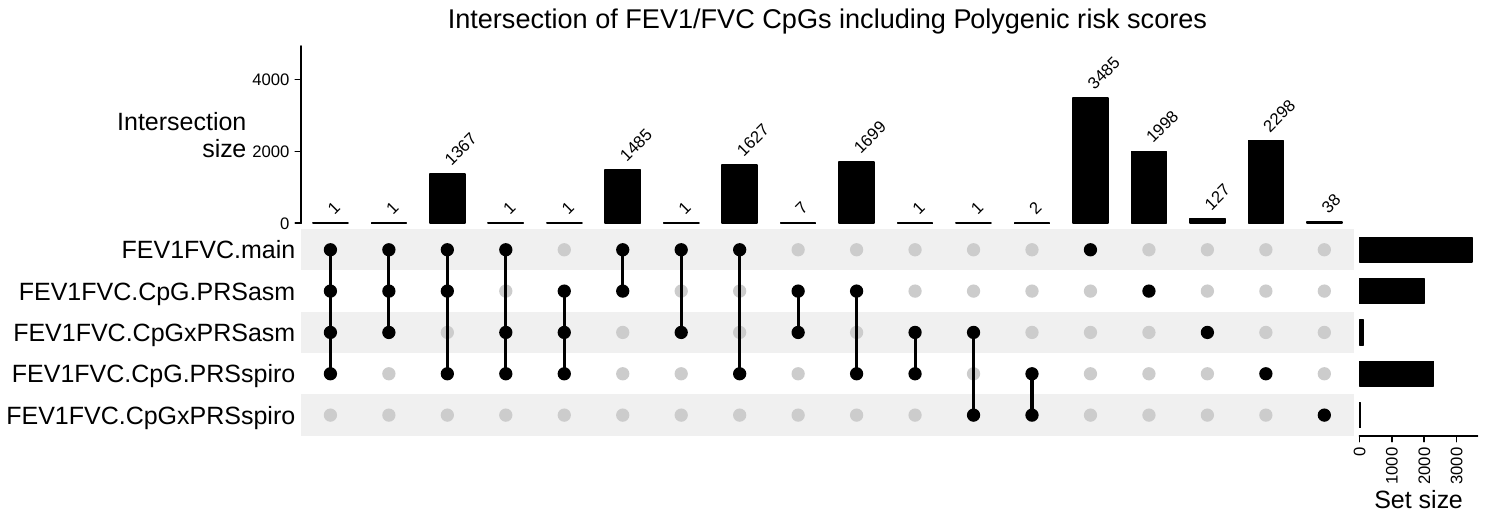

**Figure E5.** E-FORGE functional integrative analyses for DNase hypersensitivity sites, histone marks and chromatin states from the Roadmap Epigenomics Consortium using the lung function associated meta-analyzed CpGs in the same direction of effect in all the studies

A. Hypo-methylated CpGs

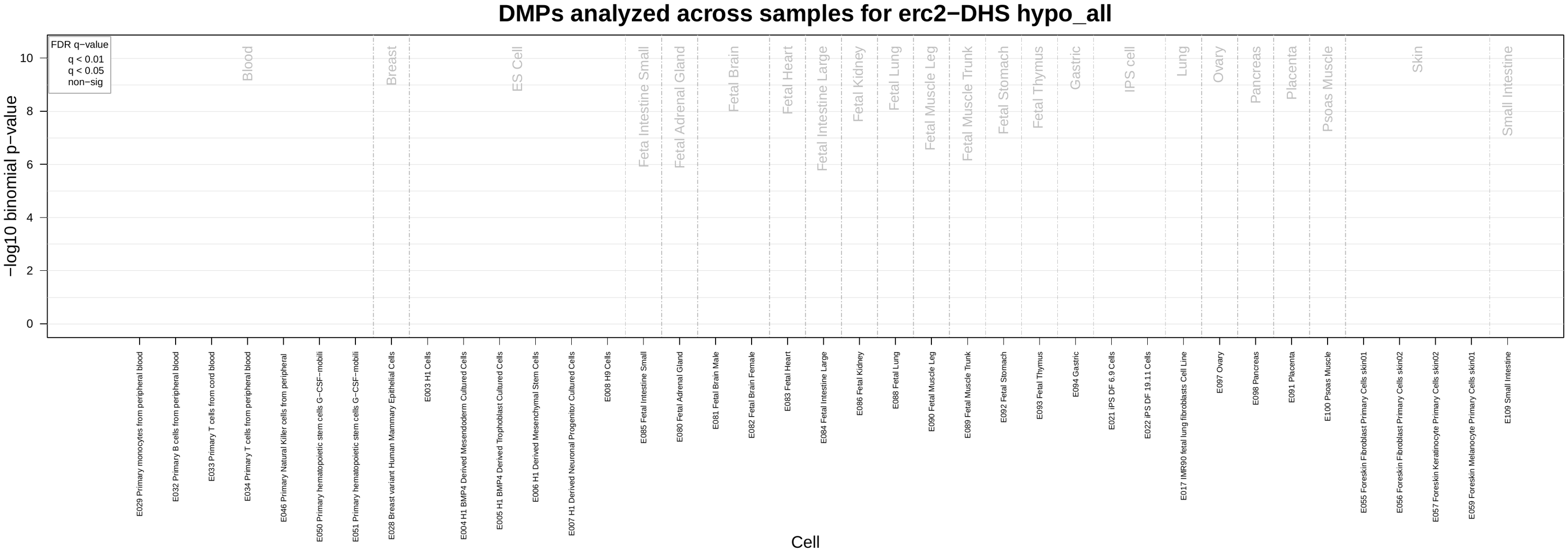

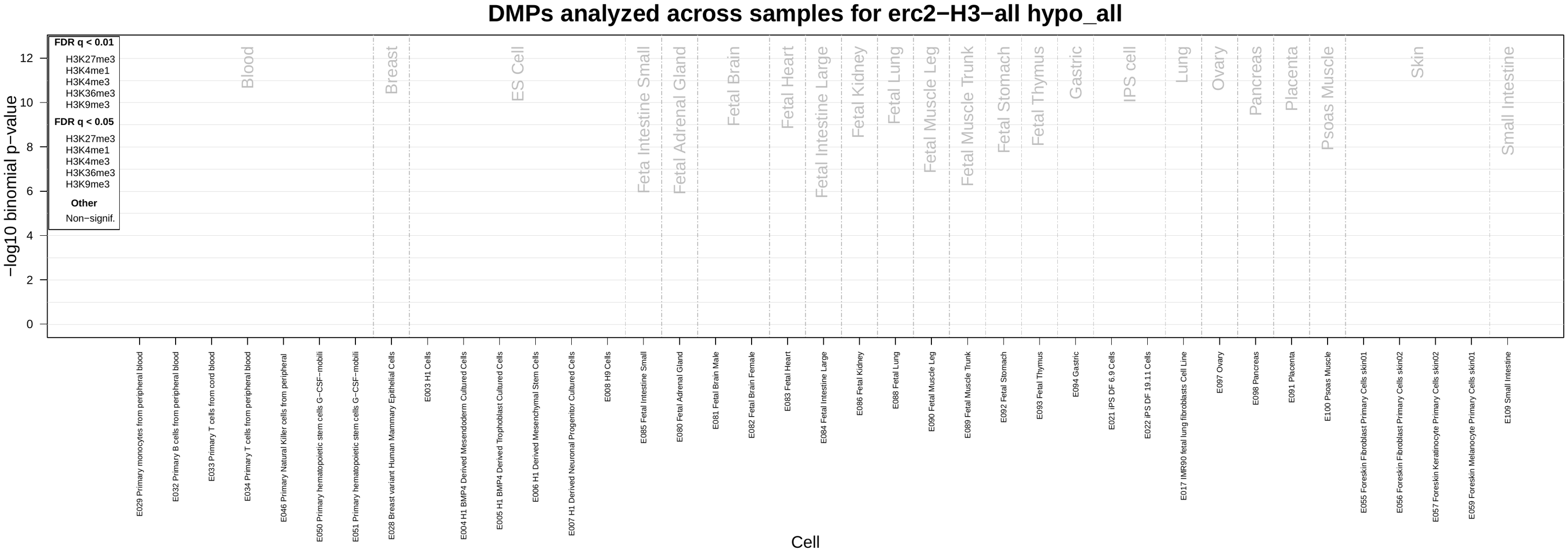

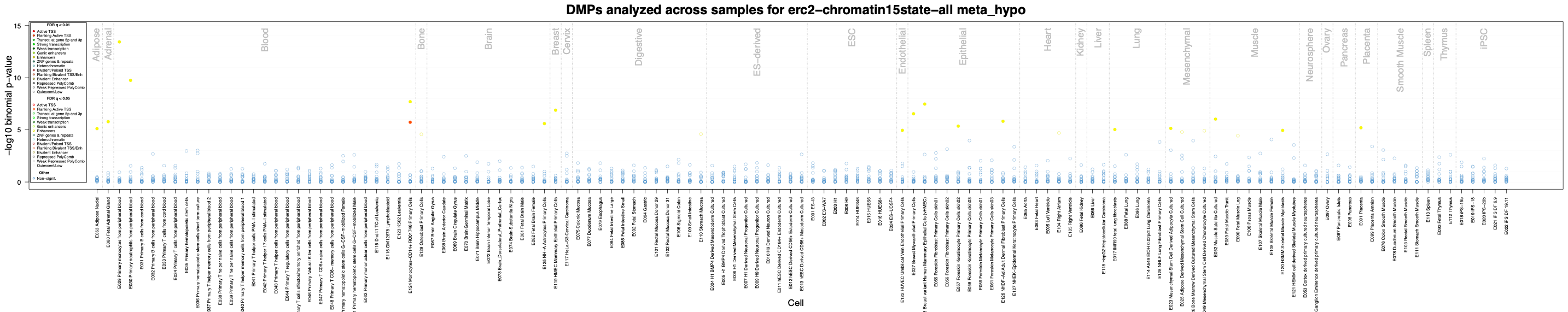

B. Hyper-methylated CpGs

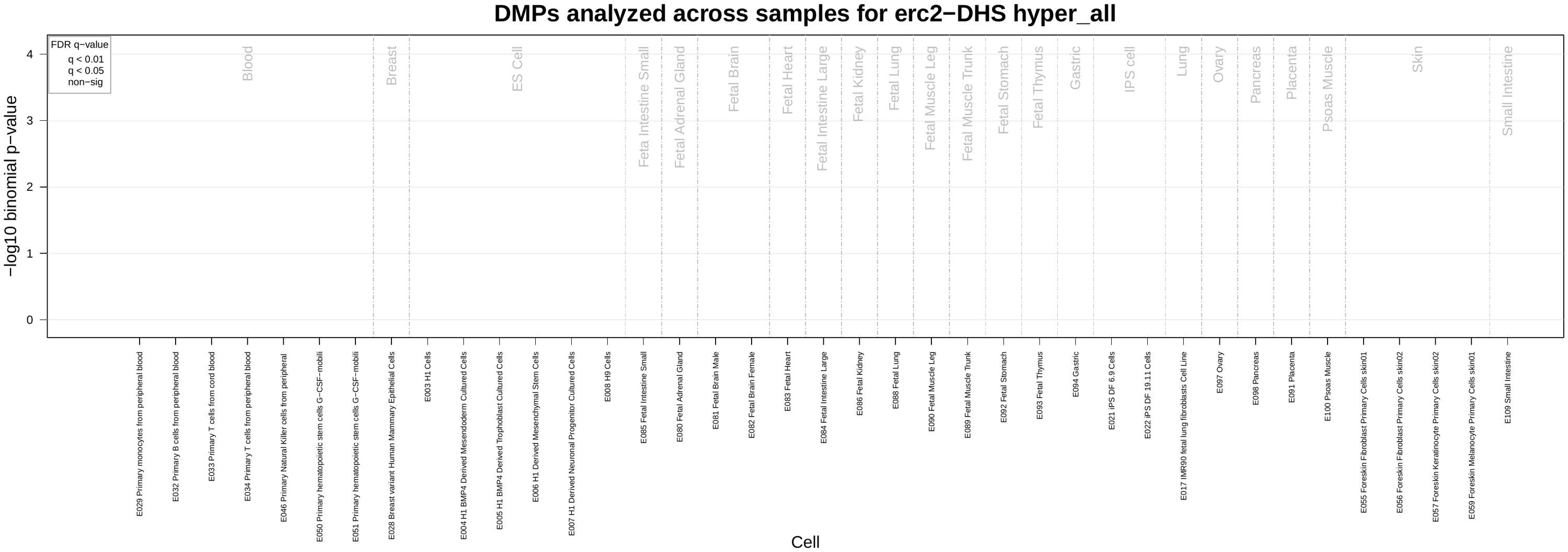

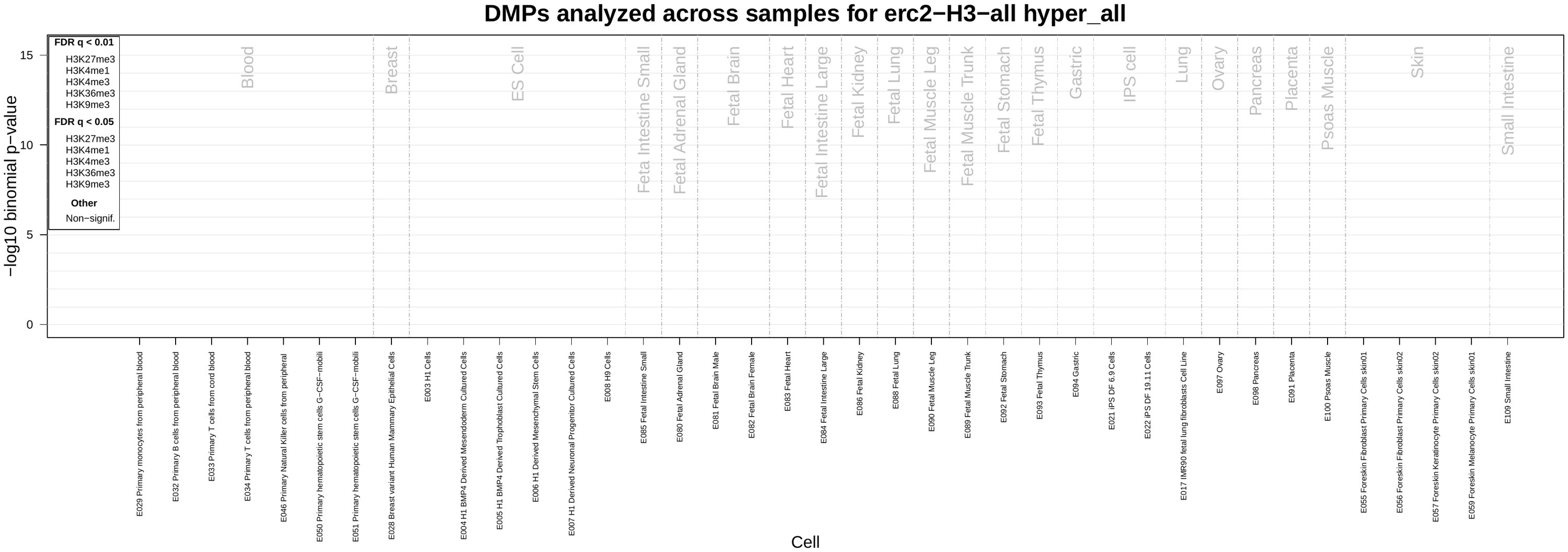

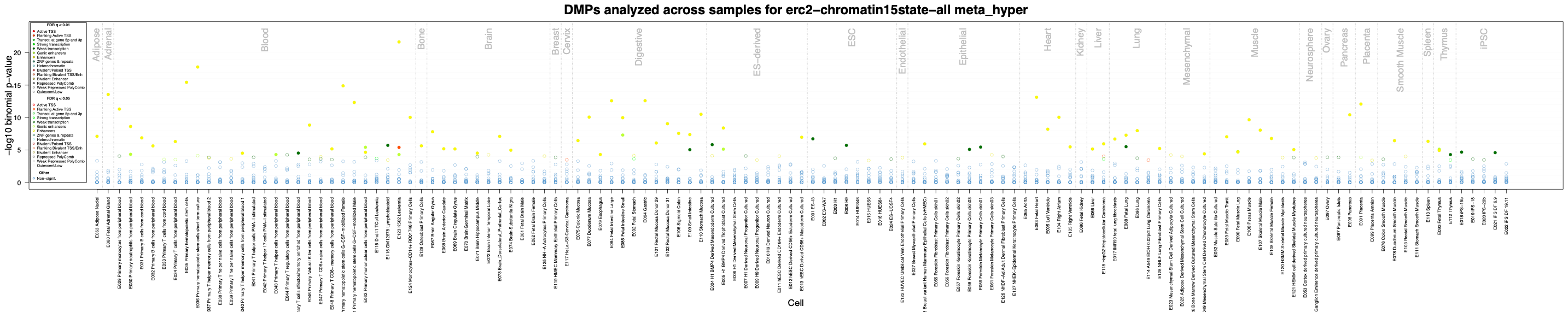

**Figure E6.** Heatmap showing the differential hub scores based on male and female specific graphical gaussian networks highlighting the sex-specific and study specific differences in hub scores across all three studies: CRA/GACRS, CAMP and VDAART. CpGs represented the nodes connected by the conditional dependence of the edges representing the direction of effect. **A.** All divergent male and female specific hub scores in the network. **B-C.** Zoomed view of the CpGs and their annotated genes with more than 20% difference in hub scores between males and females within each study population. The rows represent the CpGs and columns represent the dataset. Hub-score Key represents the range of hub scores from 0 to 1 with 0 being lowest and 1 being the highest hub score.

**A.** **B-C.**

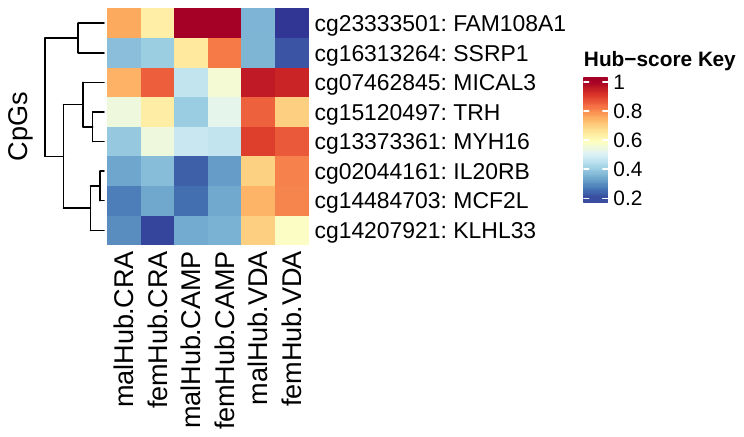

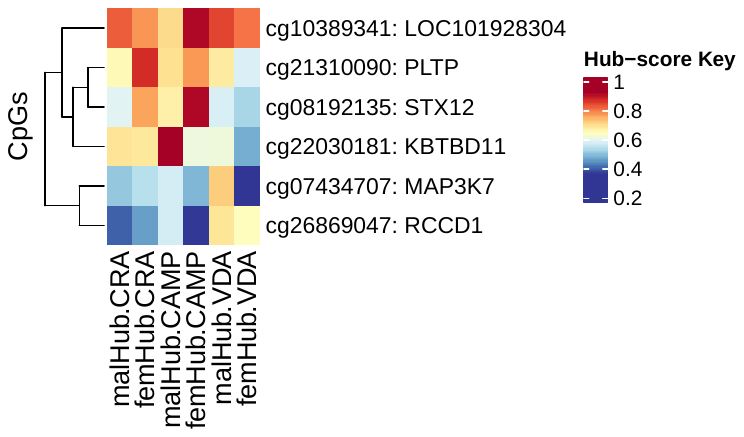

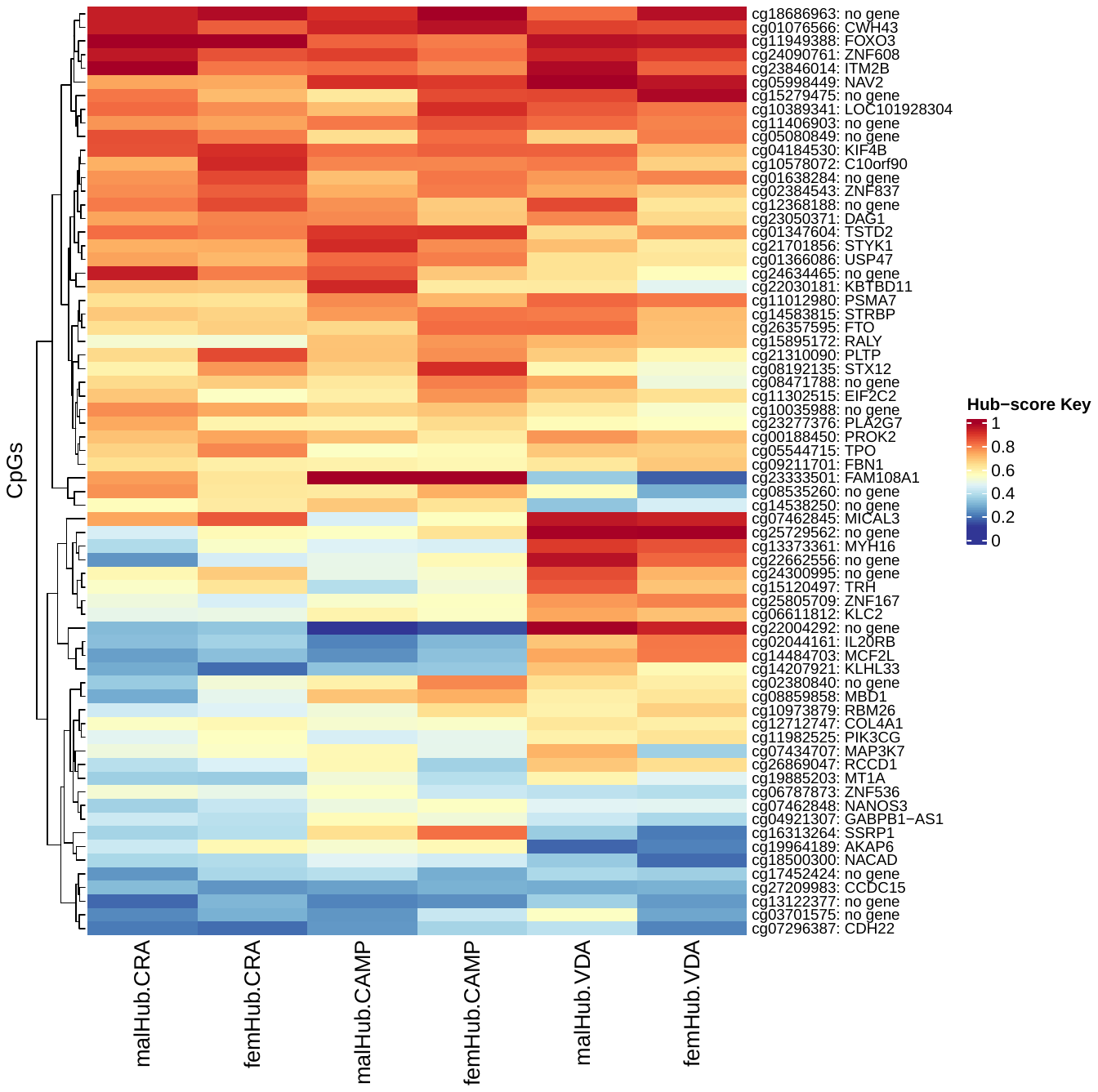
